# The proteostasis interactomes of trafficking-deficient K_V_11.1 variants associated with Long QT Syndrome and pharmacological chaperone rescue

**DOI:** 10.1101/2024.01.31.574410

**Authors:** Christian L. Egly, Lea Barny, Tri Do, Eli F. McDonald, Lars Plate, Bjorn C. Knollmann

## Abstract

**Introduction:** The voltage gated potassium ion channel K_V_11.1 plays a critical role in cardiac repolarization. Genetic variants that render Kv11.1 dysfunctional cause Long QT Syndrome (LQTS), which is associated with fatal arrhythmias. Approximately 90% of LQTS-associated variants cause intracellular protein transport (trafficking) dysfunction, which can be rescued by pharmacological chaperones like E-4031. Protein folding and trafficking decisions are regulated by chaperones, protein quality control factors, and trafficking machinery, comprising the cellular proteostasis network. Here, we test whether trafficking dysfunction is associated with alterations in the proteostasis network of pathogenic Kv11.1 variants, and whether pharmacological chaperones can normalize the proteostasis network of responsive variants.

**Methods:** We used affinity-purification coupled with tandem mass tag-based quantitative mass spectrometry to assess protein interaction changes in human embryonic kidney (HEK293) cells expressing wild-type (WT) K_V_11.1 or trafficking-deficient channel variants in the presence or absence of E-4031.

**Resultsa:** We identified 573 core K_V_11.1 protein interactors. Both variants K_V_11.1-G601S and K_V_11.1-G601S-G965* had significantly increased interactions with proteins responsible for folding, trafficking, and degradation compared to WT. We found that proteasomal degradation is a key component for K_V_11.1 degradation and that the K_V_11.1-G601S-G965* variant was more responsive to E-4031 treatment. This suggests a role in the C-terminal domain and the ER retention motif of K_V_11.1 in regulating trafficking.

**Conclusion:** Our report characterizes the proteostasis network of K_V_11.1, two trafficking deficient K_V_11.1 variants, and variants treated with a pharmacological chaperone. The identified protein interactions could be targeted therapeutically to improve K_V_11.1 trafficking and treat Long QT Syndrome.

## Introduction

The QT interval represents the duration between ventricular depolarization and repolarization in the heart measured by surface electrocardiogram. A critical component in cardiac repolarization is the voltage-gated potassium ion channel, K_V_11.1. Impairment of K_V_11.1-related ion currents (I_Kr_) can result in Long QT Syndrome (LQTS), a condition that may cause life-threatening ventricular arrhythmias. Approximately 1:2500 individuals are affected by LQTS. Of these, between 30-50% are attributed to loss of function K_V_11.1 variants, known as subtype 2 (LQT2).^1,2^

Currently, the primary treatment strategy involves beta-blockers, which target the noradrenaline system in the heart. While beta-blockers are generally effective, they do not correct underlying molecular mechanisms and fail to shorten the QT interval.^2^ Mexiletine, known for inhibiting late currents in the cardiac sodium channel (SCN5A), has demonstrated potential in shortening the QT interval in patients with LQT2 and LQT3.^3^ Despite optimal drug therapy, breakthrough cardiac events are not rare, requiring more invasive therapies such as left cardiac sympathetic denervation or the implantation of cardiac defibrillators.^2^ Thus, the development of new pharmacological treatments that directly target the underlying mechanisms of LQTS is warranted.

Around 90% of pathogenic K_V_11.1 variants cause channel dysfunction by impairing protein transport (trafficking) to the plasma membrane.^4^ However, the molecular mechanisms governing K_V_11.1 channel trafficking and protein quality control in the endoplasmic reticulum (ER) are not well understood. The K_V_11.1 protein, comprising six transmembrane domain subunits, undergoes tetramer assembly and channel folding within the ER. The channel’s pore-forming domain, located between the S5 and S6 transmembrane domains, is characterized by deep hydrophobic pockets, which binds a wide range of drugs. The resulting channel block and drug-induced long QT syndrome pose a significant challenge in drug development.^5,6^ Several known K_V_11.1 inhibitors (e.g., E-4031) can improve trafficking of some pathogenic K_V_11.1 variants.^7,8^ These inhibitors function as pharmacological chaperones by binding with high affinity to K_V_11.1, thereby stabilizing structural domains for ER export and forward trafficking.^9,10^ Although identifying drugs that improve trafficking without channel block remains a challenge, channel blocking chaperones offer the potential to elucidate the mechanisms dictating K_V_11.1 trafficking.

Here, we test the hypothesis that pathogenic K_V_11.1 variants and pharmacological chaperones that improve trafficking significantly modify protein interactions involved in folding, processing, trafficking, and degradation. These interactions constitute the proteostasis network, a diverse suite of interactions which balance a complex interplay between pro-folding and pro-trafficking factors versus pro-degradation factors, such as the ubiquitin-proteasome system, and autophagy.^11–13^ Certain K_V_11.1 protein interactions are likely crucial for proper folding and/or trafficking, while pro-degradation interactions likely lead to aberrant destruction of destabilized protein, as has been shown for other membrane proteins, such as cystic fibrosis transmembrane conductance regulator (CFTR).^14,15^ Targeting these detrimental protein interactions may hold promise to improve K_V_11.1 variant function. Multiple proteins are known to directly interact with K_V_11.1, each exerting distinct functional impacts on the channel, yet a complete understanding of the K_V_11.1 proteostasis network is lacking.^16–20^ To bridge this critical knowledge gap, we used affinity-purification coupled with mass spectrometry (AP-MS) to evaluate the proteostasis network and quantified changes in protein interactions of wild-type (WT) K_V_11.1 and two pathogenic K_V_11.1 channel variants. Such quantitative AP-MS studies have previously uncovered the proteostasis interactions of other secretory proteins, including CFTR, and quantified the alterations associated with mistrafficked disease-associated variants.^14,15,21,22^ We also tested the hypothesis that pharmacological chaperone treatment modifies the K_V_11.1 variant proteostasis networks to more closely mimic a WT state. Our work uncovered the intricate protein interaction network of K_V_11.1 and delineated unique alterations linked to two trafficking-deficient variants, and how the protein interactions are partly normalized by pharmacological chaperone treatment.

## Methods

### HEK293 cell lines

A detailed method for generation of human embryonic kidney (HEK293) cell lines expressing WT K_V_11.1, K_V_11.1-G601S, and K_V_11.1-G601S-G965* as well as cell culture conditions is previously published.^23^ The K_V_11.1-G601S-G965* cell line was specifically optimized for Tl^+^-flux assays, expresses the trafficking deficient variant G601S and removes the endoplasmic reticulum retention motif with an early truncation 17 amino acids after the amino acid G965. All HEK293 cells were cultured in HEK293 media consisting of Minimum Essential Media (MEM, Corning) containing 10% (v/v) fetal bovine serum (FBS, Gibco), and 1% Glutamax (Gibco). All cells were between passage numbers 3-30 for experiments.

### Coimmunoprecipitation (affinity purification)

HEK293 cells were cultured to full confluence in flasks ranging from 75 to 225 cm^2^. All cells were treated with either 0.1% DMSO or 10 µ M E-4031 for 24 hours. The cells were washed with Dulbecco’s phosphate buffered saline (DPBS, Gibco) without calcium and magnesium. DPBS was removed and 750 µL of ice-cold lysis buffer was added containing 50 mM Tris pH 7.5, 250 mM NaCl, 1 mM EDTA, 0.5% Igepal-CA-630 (TNI buffer) and 1x protease inhibitor cocktail (Sigma). The flasks were agitated on an orbital shaker for 30 minutes. Cell lysates were scraped from flasks into 1.7 mL Eppendorf tubes and sonicated with probe sonication (Qsonica) at 10 watts for five sequences of 3-second-bursts, separated by 7-second pause while samples were kept on ice. After sonication, samples were centrifuged at 18,000 × g for 30 minutes at 4 °C. Supernatant was collected post-centrifugation and pellets were discarded. Finally, the protein concentration was quantified using a Bicinchoninic acid assay (BCA).

Next, 10 mg of lysate representing one biological replicate was incubated for 2 hours at 4 °C with anti-Kv11.1 antibody (Cell Signaling #12889) covalently coupled to protein A/G magnetic beads (Thermo Fisher). The antibody-to-lysate ratio was maintained at 0.1 µg of antibody to 100 µg lysate. Following coimmunoprecipitation in 15 mL centrifuge tubes, complexes were transferred to 1.7 mL Eppendorf tubes. Samples were washed three times (250 µL) with a buffer containing 25mM Tris-HCl pH 7.5, 150mM NaCl, and 1mM EDTA (TN buffer). A magnetic stand was used to separate bead-Kv11.1 protein interactions from the rest of the lysate. After the final wash, all media was removed and antibody-bead/coimmunoprecipitated protein mixture was placed in a –80 °C freezer for 1 hour to facilitate later elution.

For elution, 50 µL of 0.2 M glycine (pH 2.3) buffer was added, and the sample was heated on a heating block (70 °C) for 10 minutes. Sample was collected by separating magnetic beads from eluate and then neutralized by adding 5 µL ammonium bicarbonate (1 M). Samples were stored at –80 °C until proceeding to either western blotting, total membrane staining, or liquid chromatography-tandem mass spectrometry LC-MS/MS analysis.

### Protein digestion and TMT labeling

Eluted samples were precipitated in methanol/chloroform, washed three times with methanol, and air-dried. Protein pellets were then resuspended in 4 μL 1% Rapigest SF Surfactant (Waters) followed by the addition of 10 μL of 50 mM HEPES, pH 8.0, and 32.5 μL of LCMS grade water. Samples were reduced with 5 mM tris(2-carboxyethyl)phosphine (Sigma) and alkylated with 10 mM iodoacetamide (Sigma). Next, 0.5 μg of trypsin (Sequencing Grade, Promega, or Pierce) was added and incubated for 16–18 hours at 37 °C with shaking at 700 rpm. Peptide samples were then reacted with tandem mass tag (TMT) 16/18-plex reagents (Thermo Fisher) in 40% vol/vol acetonitrile and incubated for 1 hour at room temperature. Reactions were then quenched by the addition of ammonium bicarbonate (0.4% wt/vol final concentration) and incubated for 1 hour at room temperature. TMT-labeled samples for a given mass spectrometry run were then pooled and acidified with 5% formic acid (Sigma, vol/vol). Samples were concentrated using a SpeedVac and resuspended in buffer A (95% water, 4.9% acetonitrile, and 0.1% formic acid, v/v/v). Cleaved Rapigest SF surfactant was removed by centrifugation for 30 min at 21,100 × g.

### MudPIT LC-MS/MS Analysis

LC-MS/MS data-dependent analysis (DDA) was performed using an Exploris480 mass spectrometer (Thermo Fisher) equipped with a Dionex Ultimate 3000 RSLCnano system (Thermo Fisher). Peptide samples were loaded using a high-pressure chamber onto a triphasic MudPIT column (prepared as described previously).^22^ Briefly, the MudPIT column was packed with 1.5 cm of Aqua 5 µm C18 resin (Phenomenex # 04A-4299), 1.5 cm Luna 5 µm SCX resin (Phenomenex # 04A-4398), and 1.5 cm 5 µm C18 resin. Prior to analysis, the peptide loaded MudPIT column was desalted with buffer A (95% water, 4.9% acetonitrile, 0.1% formic acid v/v/v) for 30 minutes. Peptides were eluted from the first C18 phase onto the SCX resin using a 10 µL injection of buffer A with the following 90 minute gradient: 2% B (5 minute hold) ramped to a mobile phase concentration of 40% B over 35 minutes, ramped to 80% B over 15 minutes, held at 80% B for 5 minutes, then returned to 2% B in 5 minutes and then held at 2% B for the remainder of the analysis at a constant flow rate 500 nL/min. Peptides were then sequentially eluted from the SCX stationary phase using 10 µL injections of 10, 20, 40, 60, 80, and 100% buffer C (500 mM ammonium acetate in buffer A) in buffer A. The final fraction was eluted with 10% buffer B (99.9% acetonitrile, 0.1% formic acid v/v) and 90% buffer C. Fractions were collected using the following 130-minute gradient with a constant flow rate of 500 nL/min: 2% B (6-minute hold) stepped to 5% B over 2 minutes, ramped to 35% B over 92 minutes and stepped to 85% in 6 minutes and held 85% B for 7 minutes, followed by a 17-minute hold at 2% B. For both gradients, the loading pump was held at 2% B during the duration of the analysis. During buffer pickup and loading for each method, the loading pump was held at a flow rate of 10 µL/minute for 8 mins prior to the valve switch. 5 minutes prior to the end of the analytical gradient the valve was switched to the load position to prepare for sample loading and the flow rate was adjusted from 1 µL/minute to 10 µL/minute for the remainder of the analysis. Peptides were separated using a 20 cm fused silica microcapillary column (ID 100 μm) ending with a laser-pulled tip filled with Aqua C18, 3 μm, 100 Å resin (Phenomenex # 04A-4311). Electrospray ionization was performed directly from the analytical column by applying a voltage of 2.2 kV with an MS inlet capillary temperature of 275 °C and a RF Lens of 40%.

For TMT-DDA acquisition, a 3 second duty cycle was utilized consisting of a full scan (375-1500 *m/z*, 120,000 resolution) and subsequent MS/MS spectra collected in TopSpeed acquisition mode. For MS^1^ scans, the maximum injection time was set to automatic with a normalized AGC target of 300%. Only ions with an MS^1^ intensity above 5e3 with a charge state between 2-7 were selected for MS/MS fragmentation. Additionally, a dynamic exclusion time of 45 seconds was utilized (determined from peptide elution profiles) with a mass tolerance of +/-10 ppm to maximize peptide identifications. MS/MS spectra were collected with a normalized HCD collision energy of 36, 0.7 m/z isolation window, 150 millisecond maximum injection time, a normalized AGC target of 200%, at a MS^2^ orbitrap resolution of 45,000 with a defined first mass of 110 *m/z* to ensure measurement of TMTPro reporter ions.

### Peptide Identification and Quantification

Peptide identification and TMT-based protein quantification was performed in Proteome Discoverer 2.4 (PD, Thermo Fisher). MS/MS spectra were searched using SEQUEST-HT against a Uniprot human proteome database (released 03/2014 and containing 20,337 entries; supplemented with common MS contaminants and curated to remove redundant proteins/ splice isoforms) and a decoy database of reversed peptide sequences. Searches were carried out according to the following parameters: 20 ppm peptide precursor tolerance, 0.02 Da fragment mass tolerance, minimum peptide length of 6 AAs, trypsin cleavage with a maximum of two missed cleavages, dynamic modifications of Met oxidation (+ 15.995 Da), protein N-terminal Met loss (-131.040 Da), protein N-terminal acetylation (+ 42.011 Da), protein N-terminal Met loss and acetylation (+ 89.030), and static modifications of Cys carbamidomethylation (+ 57.021 Da) and TMTPro (N-terminus/ Lys, + 304.207 Da). IDs were filtered using Percolator with a peptide level FDR of 1%. The Reporter Ion Quantification node in PD was used to quantify TMT reporter ion intensities. Only reporter ions with an average signal-to-noise ratio (S/N) greater than 10 to 1 (10:1) and a percent co-isolation less than 25 were utilized for quantification. TMT reporter ion intensities were summed for peptides belonging to the same protein, including razor peptides. Proteins were filtered at 1% FDR and protein grouping was performed according to the parsimony principle. Protein identifications and TMT quantification can be found in **Supplemental Table 1**.

### Proteomics analysis

Protein abundances for all channels within each mass spectrometry run were median normalized (N = 4 runs). The median normalized quantifications are included in **Supplemental Table 1.** To determine K_V_11.1 enriched interactions, data were log_2_ normalized and each protein pulled down with K_V_11.1 was compared to the corresponding log_2_ TMT intensity for each protein pulled down in untransfected cells. Significance was determined using multiple t-tests with false rate discovery correction by Benjamini, Krieger, and Yekutieli (q < 0.05). For subsequent analysis, we scaled log_2_ protein abundances to the K_V_11.1 protein within that channel. We then scaled the log_2_ protein abundances to the respective protein in WT K_V_11.1 samples from the same mass spectrometry run for a common reference point. The individual replicates from separate runs were compiled and averaged to result in the log_2_ fold change for each protein compared to quantified interaction in WT K_V_11.1 samples (log_2_ fold change/WT). We only included proteins identified and quantified in both mutant variants from the mass spectrometry runs.

### Covalent Antibody-bead crosslinking

This procedure was adapted from Pankow, et al.^24^ Pierce Protein A/G Magnetic Beads (ThermoFisher) containing a recombinant Protein A/G (∼50,500 kDa) that combines the IgG binding domain of both Protein A and Protein G were used to cross-link anti-Kv11.1 antibodies (Cell Signaling). Stock beads with a concentration of 10 mg/ml were washed four times with 10 volumes of Dulbecco’s Phosphate Buffered Saline (DPBS, ThermoFisher). Excess DPBS was removed, followed by the addition of an appropriate amount of anti-Kv11.1 antibodies to the beads to make the final concentration antibody/bead of 60 µg/100 µL, which is equivalent to 42.8 µL of antibody per 100 µL of stock beads. Then, the antibody and beads were gently mixed for two hours at room temperature (20-25°C) on a rotator to allow the binding of the antibody to the beads. Next, the beads were pulled with a magnetic stand and washed twice with 1 mL of freshly made 100 mM sodium borate (pH 9.0, Sigma-Aldrich). To start the cross-linking reaction, beads were resuspended in 1 mL of 100 mM sodium borate (pH 9.0) followed by the addition of dimethylpimelimidate powder (DMP, ThermoFisher) to a final DMP concentration of 25 mM. This is done as cross-linking is more efficient at pH levels greater than 8.^25^ The bead-antibody mixture was rotated for 30 minutes at room temperature on a rotator, then the beads were pulled with a magnetic stand and the supernatant was discarded. After that, beads were washed once with 1 mL of 200 mM newly prepared ethanolamine (pH 8.0) (Sigma-Aldrich), resuspended in 1 mL of 200 mM ethanolamine, and incubated for 2 hours at room temperature with gentle mixing. Finally, beads were washed five times with 1 mL of DPBS to remove excess ethanolamine and stored in DPBS with 0.1% Tween-20 (v/v) (Sigma-Aldrich) and 0.02% sodium azide (wt/vol) (Sigma-Aldrich).

### Gene Ontology Analysis

Biological processes of K_V_11.1 protein interactors were analyzed using DAVID.^26,27^ Gene symbols were converted to ENTREZ gene IDs and submitted in list format. Enriched gene ontology terms were picked from the biological process direct terms with false discovery rate correction with adaptive linear step-up adjusted p-values as discussed in Benjamini and Hochberg (q < 0.05).^28^

### Immunoblots (trafficking efficiency)

Increased K_V_11.1 variant trafficking is observed on a western blot by increased proportion of trafficked, fully glycosylated (∼155 kD) protein compared to the untrafficked, core glycosylated (∼135 kD) protein. HEK293 cells were cultured in 6-well plates in the presence of vehicle (0.1% DMSO) or E-4031 (10 µM) for 24 hours at 37 °C and 5% CO_2._ Cells were lifted using TrypLE express (ThermoFisher), centrifuged, pelleted, and lysed with 100 µL TNI buffer for 30 minutes at 4 °C. Samples were centrifuged at 18,000 × g (4 °C) for 30 minutes to remove insoluble material. The supernatant was then quantified using BCA reagent (ThermoFisher).

15 µg of protein were loaded into individual wells of a 4-20% Tris-Glycine eXtended (BioRad) precast gradient gel and run at 50 V for 30 minutes to resolve the stacking layer, then 150 V for 1.5 hours. The gel was transferred to a 0.45 µm PVDF membrane and blocked for 1 hour with 5% (wt/vol) molecular-grade non-fat milk. Primary anti-K_V_11.1 antibody (Cell Signaling, 1:2000) was incubated 16 hours at 4 ⁰C. Then membranes were incubated with secondary anti-rabbit IgG horse radish peroxidase (Promega, 1:5000) for 1 hour. Membranes were incubated for 1 minute using SuperSignal West Pico PLUS Chemiluminescent Substrate (ThermoFisher) and imaged with an iBright 1500 (ThermoFisher).

### Patch Clamp Electrophysiology (K_V_11.1 function)

Patch clamp electrophysiology of HEK293 cells was performed as previously described.^23^ K_V_11.1 current was measured in HEK293 cells by whole-cell, patch-clamp electrophysiology using a Multiclamp 700A amplifier (Axon Instruments), Digidata 1322A analog-to-digital converter (Axon Instruments), and TE200 microscope (Nikon). A total of 15,000 cells were plated on 35 mm dishes with 20 mm glass-bottom wells (Cellvis D35-20-1.5-N). Cells were cultured a minimum of two days in a 37 ⁰C incubator with 5% CO_2_. Cells were treated with either vehicle (0.1% DMSO) or E-4031 (10 µM) for 24-hours before experiments. External patch-clamp solution contained (in mmol/L) NaCl 137, KCl 4, CaCl_2_ 1.8, MgCl_2_ 1, Glucose 10, and HEPES 10 (pH 7.4 using NaOH). Internal solution contained (in mmol/L) KCl 137, MgCl_2_ 1, EGTA 5, HEPES 10, MgATP 5 (pH 7.2 using KOH). Glass capillaries (World Precision Instruments) were pulled to 1.5-3.5 megaohm internal resistance. Prior to recordings, cells were perfused with external patch solution (2.5-4 mL/minute) for 30 minutes at 30 ⁰C to remove residual drug. Recordings from single, isolated cells were performed at room temperature (∼23-24 ⁰C).

### Statistical analysis

Where appropriate unpaired student’s t-test, ANOVA with post-hoc Tukeys multiple comparisons, and Kruskal-Wallis with post-hoc Dunn’s multiple comparisons were performed. To assess whether data were parametric or not, we used the Shapiro-Wilk test for data with multiple samples. Outliers were removed from patch clamp electrophysiology data using the Robust regression and Outlier Removal (ROUT) test with Q = 5%. All statistical tests were performed in GraphPad Prism

## Results

### Coimmunoprecipitation of K_V_11.1 and two trafficking-deficient variants

Our primary objective was to understand how the proteostasis network differs between WT K_V_11.1 and pathogenic K_V_11.1 variants with impaired trafficking. We utilized a well-documented trafficking-deficient variant, K_V_11.1-G601S, and a double mutant with the same variant and a truncation (K_V_11.1-G601S-G965*). The double mutant lacks the endoplasmic reticulum retention signal RXR (amino acids 1005-1007) and is also trafficking-deficient due to the G601S mutation. Henceforth, we will denote this variant as G965*. The G965* variant demonstrated enhanced responsiveness to E-4031 chaperone treatment in functional assays, making it a more suitable cell line to develop a high-throughput drug screen.^23,29^ This finding suggested that the proteostasis network of the truncated variant may differ from the full length G601S variant.

To map the complete interactome of the K_V_11.1 protein, including all direct and indirect protein interactions, we employed an affinity purification approach coupled with tandem mass tag (TMT) labeled mass spectrometry on whole-cell lysates from HEK293 cells **(Fig. 1A)**. In this procedure, cells are lysed gently to preserve native interactions, co-immunoprecipitated using anti-K_V_11.1 antibody, eluted, and digested. Subsequently, peptides from each sample are labeled with an isobaric TMT reagent, allowing for simultaneous recordings of protein abundances in up to 18 samples in a single mass spectrometry run. Differences in protein contacts and interactions between WT ion channels and their genetic variants are then quantifiable. While these networks remain largely unknown for K_V_11.1, research has illuminated the networks for several other ion channel disorders.^14,15,30,31^

**Figure 1:**
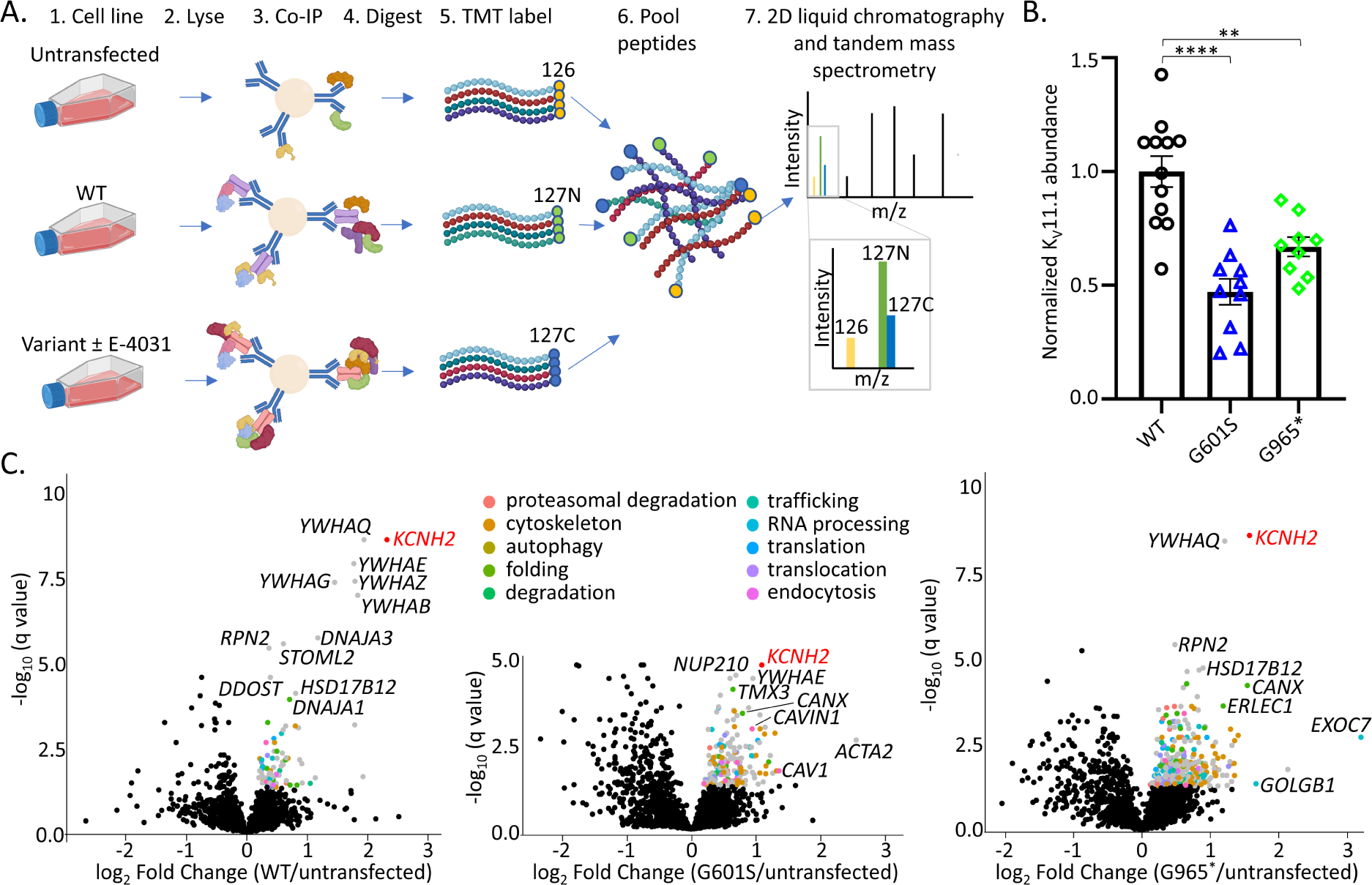
Identification of K_V_11.1 protein interactors. **(A)** Multiplexed affinity-purification mass spectrometry workflow for mapping K_V_11.1 and variant protein interactions. **(B)** Bar graph illustrating the normalized abundance of K_V_11.1 in TMT-labeled channels from replicates, with bars representing mean ± SD. **(C)** Volcano plots depicting K_V_11.1 interactors identified via multiplexed AP-MS proteomics in HEK293 cells expressing WT K_V_11.1 and two trafficking-deficient variants. The plots highlight significantly enriched proteins (shown in gray, q < 0.05 by multiple t-tests with false discovery rate correction using two-stage linear step-up of Benjamini, Krieger, and Yekutieli), further divided using manual gene ontology terms (N=10) and color-coded for clarity. Key proteins like K_V_11.1 (*KCNH2,* red) and other genes are labeled.

To determine the proteostasis network of WT K_V_11.1, two trafficking-deficient variants, and variants treated with the pharmacological chaperone E-4031, we conducted co-immunoprecipitations (Co-IP) and independent mass spectrometry runs for multiple replicates per mutation and treatment (WT, N=12; G601S, N=10; G601S E-4031, N=6; G965*, N=9; G965* E-4031, N=8) (**Supplementary Table 2**). We included untransfected mock controls (N=12) to assess the abundance of background proteins during the Co-IP procedure.

The K_v_11.1 bait protein was confidently identified in all runs (mean ± SD, 48.8 ± 4.2 % sequence coverage). Quantification of the TMT reporter ion intensities showed that the total abundance of WT K_V_11.1 protein measured by mass spectrometry was elevated compared to the two K_V_11.1 variants **(Fig. 1B)**, suggesting the K_V_11.1 variants undergo increased protein degradation, as has been reported for other ion channels with trafficking-deficiency.^14^ We next defined statistically significant interactors by comparing the log_2_ fold enrichment in the Kv11.1 Co-IP conditions compared to untransfected sample. We conducted a median normalization based on total protein abundances for each TMT channel to account for slight variations in the overall Co-IP enrichment (**Fig. S1**). We filtered interactors based on a positive log2 fold change and multiple t-tests with false discovery rate of q < 0.05 (**Supplementary Table 3**). In cells expressing WT K_V_11.1, we identified 88 significantly enriched protein interactions, 192 in the G601S variant, and 367 in the G965* variant (**Fig. 1A, Fig. S2A-B**). These data suggest that trafficking-deficient variants have an enriched set of protein interactors, which is consistent with their increased surveillance by the proteostasis network (**Fig. S2C**). The G965* variant shares 50 interactors with WT while the G601S variant only shares 32 interactors, indicating that G965* interactome more closely resembles that of WT Kv11.1 (**Fig. S2C-F**).

We then compiled the interactors from each condition into a master list of interactors based on significance in at least one condition compared to untransfected cells. This resulted in 573 enriched K_V_11.1 protein interactions. We identified several known K_V_11.1 protein interactions such as 14-3-3ε (*YWHAE*) and calnexin (*CANX*) **(Fig. 1C)**.^9,32–34^ Using manual annotation from Uniprot, we classified each protein interaction with a unique gene ontology (GO) term to represent the most prominent biological process (**Supplementary Table 4**).

### Cellular model of K_V_11.1 trafficking to plasma membrane

Since a global understanding of K_V_11.1 interactions remained unknown until this study, we utilized the Database for Annotation, Visualization, and Integrated Discovery (DAVID) bioinformatics to guide gene ontology (GO) term classifications and cellular localization of protein interactors. Utilizing this database, we developed a cellular model for K_V_11.1 biogenesis **(Fig. 2)**. Significant, intracellular protein interactions with K_V_11.1 were found from the nucleus (*NUP85/88/93/*107/155/160/210/214) to the plasma membrane (*CAV1* and *CAVIN1*). We found several significant interactions responsible for mRNA translation, which include eukaryotic initiation factors (*EIF2/3/4*) and ribosomal protein subunits (*RPS3/5*). These data are consistent with reports that voltage gated potassium channels associate with ribosomal proteins and tetramer formation can occur prior to polypeptide exit from the ER translocation channel.^35^ We identified numerous folding chaperones that include components of the heat shock 40 complex (*DNAJA1/2/3*), a peptide prolyl isomerase (FK506 binding protein 8, *FKBP8)*, the coiled-coil domain containing 47 protein (*CCDC47)*, calmegin (*CLGN*), and lectin-based chaperones calnexin (*CANX)*, and the endoplasmic reticulum lectin 1 (*ERLEC1)*. Calnexin reportedly interacts with K_V_11.1, which is significantly reduced after pharmacological chaperone treatment of the trafficking-deficient N470D variant.^9^ DNAJB12 has been shown to be responsible for K^+^ channel tetrameric assembly in a heat shock 70 independent manner.^36^ Several proteins involved in assisting with proteasomal degradation of unfolded proteins included the transitional ER ATPase p97 (*VCP*), the ER lipid raft-associated proteins 1&2 (*ERLIN1/2*), the E3 ubiquitin-protein ligase itchy homolog (*ITCH*), and a core component of the multiple cullin-RING E3 ligases (cullin-3, *CUL3*). Proteins responsible for antegrade (ER to Golgi) and retrograde (Golgi to ER) trafficking included multiple vesicle coating proteins (*COPA*, *COPB1/2*, and *COPG2*), vesicle-associated membrane proteins (*VAPA* and *VAPB*), vesicle trafficking protein (*SEC22B*), ER lumen protein retaining receptor 1 and 2 (*KDELR1* and *KDELR2*), the transmembrane emp24 domain-containing protein 9 (*TMED9*), and the ADP ribosylation factor 4 (*ARF4*). Plasma membrane transport protein interactions included filamin A (*FLNA*), plakophilin 2 and 3 (*PKP2/3*), Hyccin (*HYCC1*), and myosin regulatory light chain 12A (*MYL12A*). Proteins responsible for endocytosis and lysosomal degradation included caveolin-1 (CAV1), caveolae associated protein (*CAVIN1*), protein spire homolog 1 (*SPIRE1*), the phosphatidylinositol-binding clathrin assembly protein (*PICALM*), multiple Ras-related proteins (*RAB10*, *RAB32*, and *RAB14*) and the spectrin beta chain, non-erythrocytic 2 protein (*SPTBN2*). While not comprehensive, these proteins form key components of the K_V_11.1 interactome.

**Figure 2:**
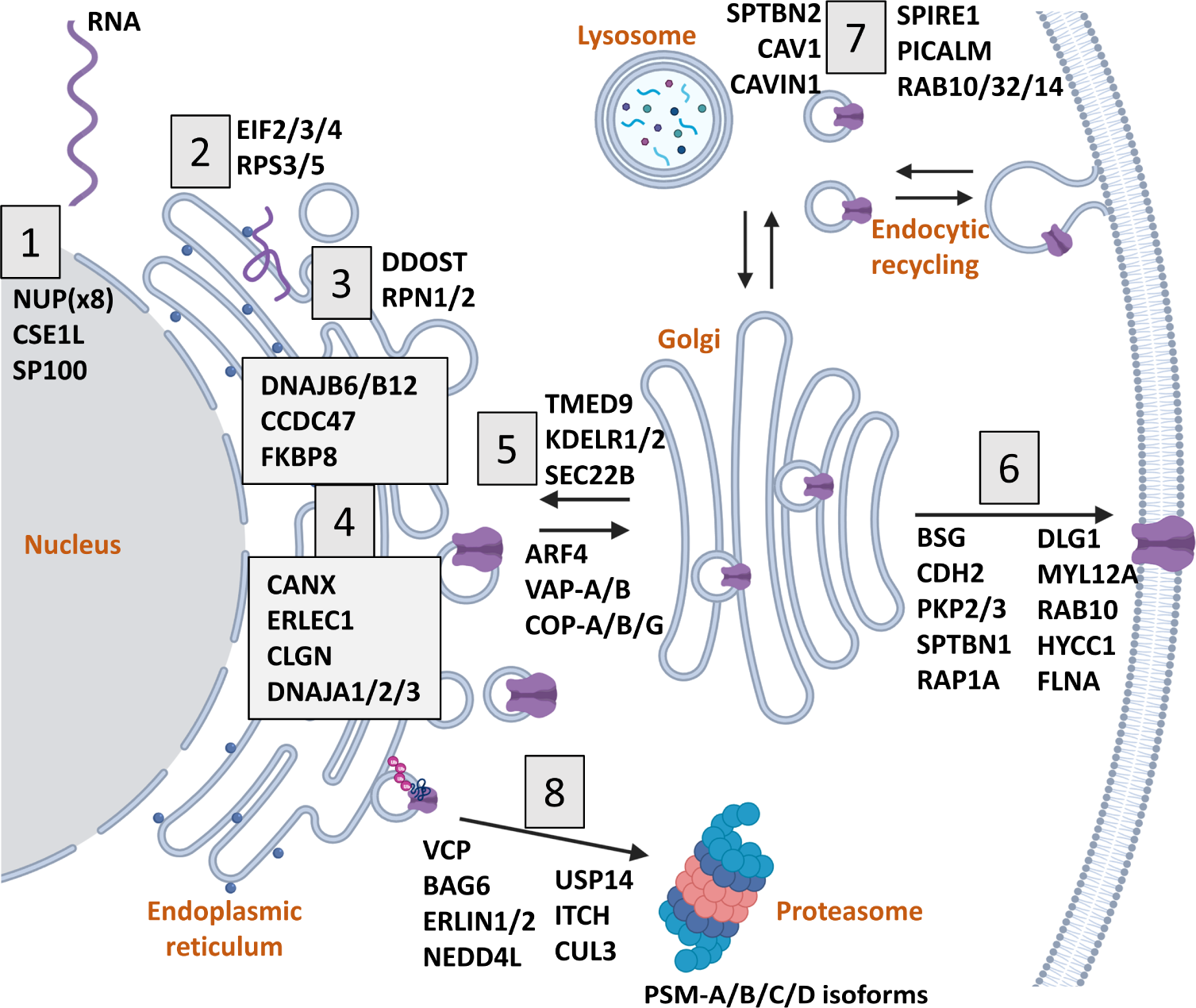
Diagram of K_V_11.1 Protein Trafficking and Interaction Networks. The cellular model highlights K_V_11.1 protein interactions involved in 1) Nuclear import/export, 2) Translation, 3) Early N-linked glycosylation, 4) Folding chaperones in the endoplasmic reticulum, 5) ER-Golgi transport, 6) Plasma membrane transport, 7) Endosomal recycling, and lysosomal degradation, and 8) Proteasomal degradation. Each cellular organelle or critical process is marked in orange.

### Trafficking-deficient variants show increased protein interactions

To quantitate variations in protein interactions between WT and variants, it was important to account for any variation in K_V_11.1 bait protein intensities (**Fig. 1B**). Therefore, the TMT abundances for each interactor from the master list were normalized to the K_V_11.1 bait protein abundance. We then used the WT K_V_11.1 conditions as a common reference point in each run. For each interactor, we calculated the log2 enrichment fold change comparing each variant and/or treatment condition to WT (**Supplementary Table 5**). This established the enrichment fold change for each interactor from K_V_11.1 variants relative to the same protein in WT samples.

In aggregate, both trafficking-deficient K_V_11.1 variants exhibited a significant increase in protein interactions compared to WT, indicating either more extensive interaction networks within the ER and other cellular regions or prolonged interactions with quality control factors **(Fig. 3A)**. Across all 573 enriched protein interactions, G601S had an elevated average Log_2_ fold change from WT compared to the G965* variant (G601S 1.69±0.02 vs. G965* 1.04±0.02, mean ± SEM). We utilized DAVID to determine significantly enriched pathways from the input of 573 enriched K_V_11.1 interacting proteins.^26^ Manual GO term assignment categorized each protein to one dominant function, whereas DAVID annotations group single proteins into all associated terms. The top four enriched DAVID GO terms from all 573 K_V_11.1 protein interactors included actin cytoskeleton and filament organization, proteasomal degradation, and mRNA stability **(Fig. 3B).** We further examined 10 enriched GO terms responsible for trafficking and degradation as defined by DAVID to detail specific changes in protein interactions across various folding and trafficking related domains **(Fig. 3C & Supplemental Table 6)**. These interactions represent unique functions in the cells such as mRNA translation initiation, ER associated protein folding, ER to Golgi vesicle or non-vesicle mediated transport, other transport specific to localization to the plasma membrane, endosomal recycling (vesicle-mediated transport), overall protein degradation via ER-associated degradation or proteasome specific degradation, and actin nucleation, which plays a variety of roles in maintaining the actin cytoskeleton. We discovered several proteins consistently enriched across all conditions, indicating high-confidence interactions. These included proteins involved in early N-linked glycosylation and degradation: ribophorin II (*RPN2*) and dolichyl-diphosphooligosaccharide-protein glycosyltransferase subunit (*DDOST*), folding glucosidase II alpha subunit (*GANAB*) and calnexin (*CANX*), and neural precursor cell expressed developmentally down-regulated protein 4-like (*NEDD4L*) **(Fig. 3D-H)**. Notably, we also identified a protein interaction between K_V_11.1 and the very-long-chain 3-oxoacyl-CoA reductase (*HSD17B12*), which is responsible for fatty acid synthesis and involved in converting estrone to estradiol **(Fig. 3I)**. Interestingly, G965* interactions levels were significantly decreased compared to G601S for interactors *NEDD4L*, *RPN2*, *DDOST*, and *GANAB* **(Fig. 3D-I)**.

**Figure 3:**
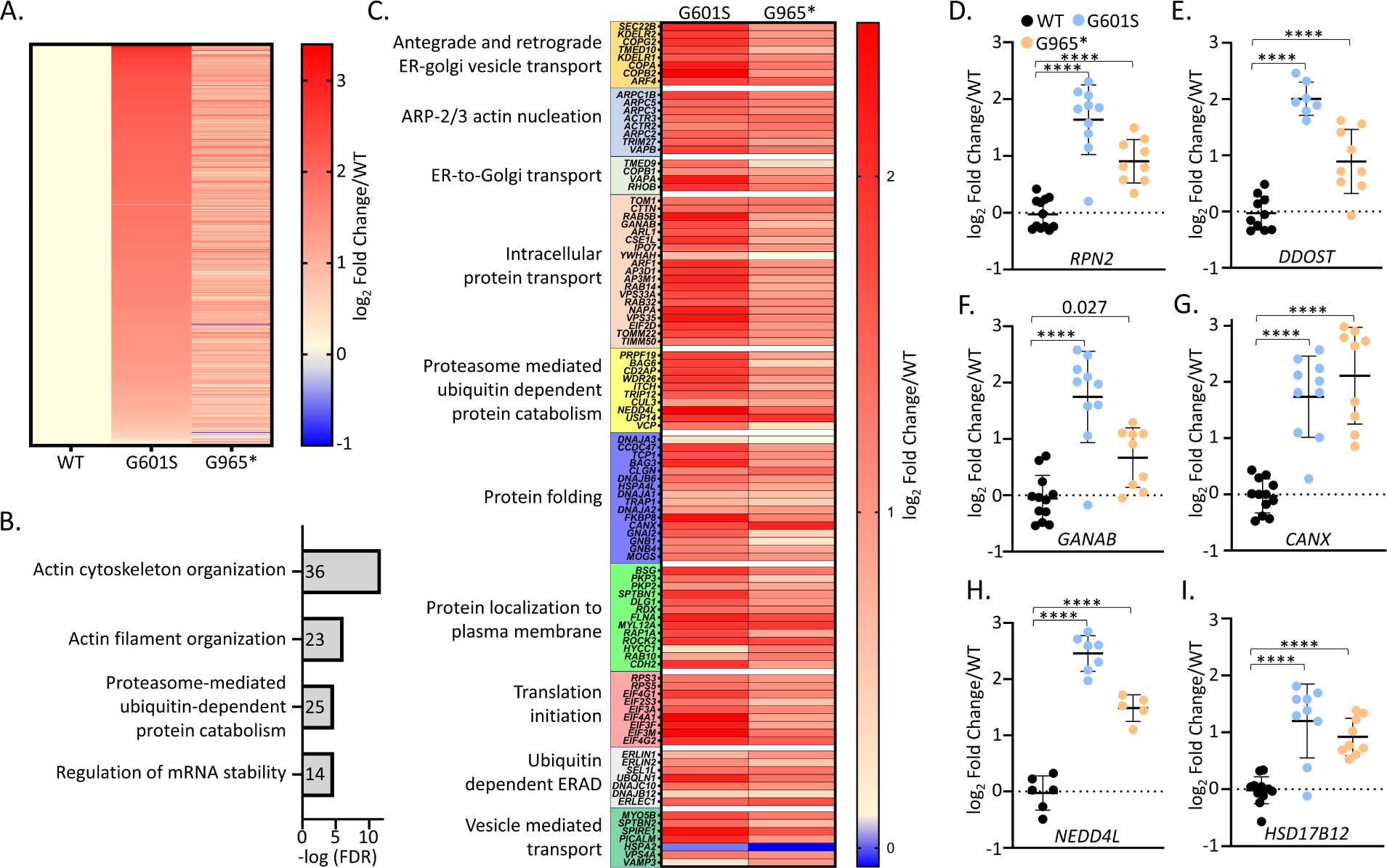
Comparative protein interaction dynamics in K_V_11.1 and variants. **(A)** Heatmap illustrating the 573 enriched protein interactions from K_V_11.1 AP-MS experiments. Log_2_ fold changes are scaled to WT treated with 0.1% DMSO and ranked from highest to lowest fold change for G601S. **(B)** The top four enriched pathway terms identified by DAVID analysis for the 573 enriched K_V_11.1 protein interactors, with the count of associated proteins for each GO term displayed within the bars. **(C)** Heatmap representing log_2_ fold change from WT K_V_11.1 for select interactions, categorized and assessed using DAVID clustering to illustrate their potential functional significance and interconnectivity. **(D-I)** Dot plots illustrating quantified protein enrichment for individual genes, each dot representing a distinct biological replicate. In heat maps and dot plots, log_2_ fold change is scaled to WT K_V_11.1. Statistical significance was calculated using one-way ANOVA with post-hoc Tukey’s multiple comparisons test and p values are depicted by **** < 0.0001, *** = 0.0001. Bars represent mean ± SD.

To assess the interaction changes between WT and the two mutant variants in an unbiased fashion, we carried out hierarchical clustering to identify groups of interactors that exhibit common patterns (**Fig. 4A and Fig. S4**). Hierarchical clustering revealed 6 clusters containing the most proteins in cluster 1, 3, and 5 (127, 152, and 108 proteins each respectively), with fewer proteins in clusters 2, 4, and 6 (49, 74, and 62 respectively). Among these clusters, we identified two clusters (4&6) that displayed similarly elevated interactions for the two variants when compared to WT. The other four clusters had elevated interactions for G601S compared to G965*, albeit the magnitude differed (**Fig. 4B**). To determine pathways overrepresented in each cluster, we carried out DAVID analysis to show which biological processes were enriched. For cluster 4 with similar interactions for the two mutants, we identified an enrichment of YWHA proteins responsible for protein targeting, positive regulation of protein insertion into mitochondrial membrane involved in apoptotic signaling pathway, and cellular protein localization (**Fig. 4C**). Cluster 6 was enriched for actin cytoskeleton organization (**Fig. 4D**). All four clusters that had divergent interactions between the two variants were enriched for proteasomal mediated ubiquitin dependent protein catabolism in cluster 1 (**Fig. 4E**), proteasomal catabolic processes in cluster 2 (**Fig. 4F**), ARP-2/3 actin nucleation in cluster 3 (**Fig. 4G**), and actin filament bundle assembly and network formation in cluster 5 (**Fig. 4H**). The data indicate a substantial portion of protein interactions involving K_V_11.1 occurs within the actin cytoskeleton, which was enriched in the largest clusters 3 and 5 as well as cluster 6. Notably, many of these interactions differ between the variants (3 and 5) while some are similar between the two (cluster 6). Proteasomal mediated degradation emerged as the dominant degradation pathway for K_V_11.1, featured in clusters 1 and 2.

**Figure 4:**
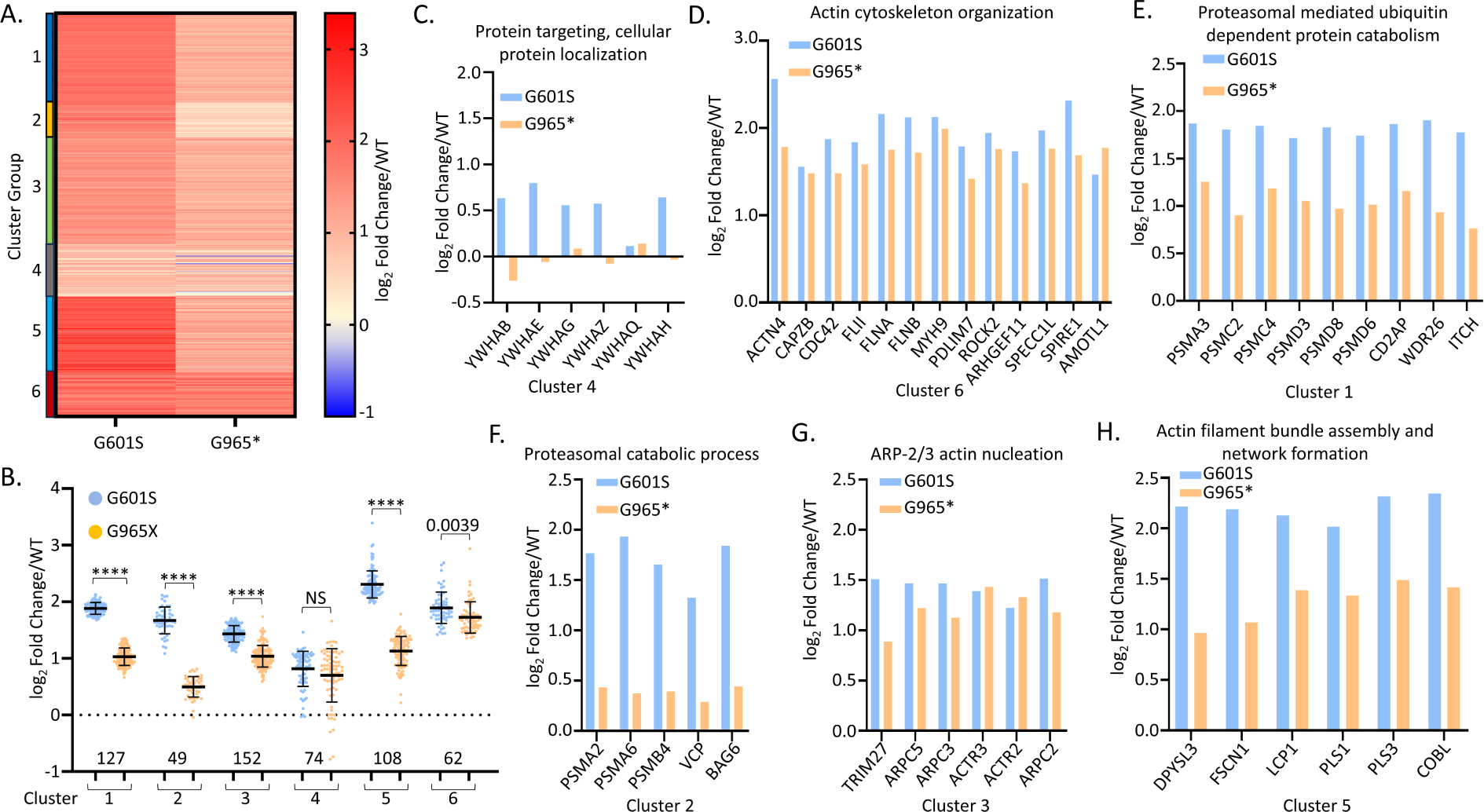
Cluster Analysis of trafficking-deficient K_V_11.1 interactomes. **(A)** Hierarchical clustering of 573 protein interactors arranged by intensity similarity. Abundances for G601S and G965* variants in relation to WT K_V_11.1 were transformed into a Euclidian distance matrix and clustered using Ward’s minimum variance method. Each row represents an interactor with color indicating log_2_ fold change normalized to WT. **(B)** Swarm plots for all clusters depicting the log_2_ fold change normalized to WT for each variant and cluster. Statistical significance was calculated by ANOVA with post-hoc Tukey’s multiple comparisons test and p values are depicted by **** < 0.0001. Bars represent mean ± SD. (**C-H**) Bar charts of individual protein comparisons selected by enriched protein GO terms after DAVID analysis. All samples were treated with vehicle (0.1%) or E-4031 (10 µ M) for 24-hours prior to sample collection.

### K_V_11.1 proteostasis network changes in response to pharmacological chaperone E-4031

We next explored how chaperone treatment with E-4031 alters interactions of trafficking-deficient variants. The potent K_V_11.1 channel inhibitor, E-4031, a methanesulfonamide class III antiarrhythmic similar to dofetilide and ibutilide, significantly enhances membrane trafficking of K_V_11.1 variants but not WT **(Fig. 5A&B)**. Next, we carried out functional measurements of channel conductance. Treatment with E-4031 for 24-hours significantly enhanced K_V_11.1 variant currents **(Fig. 5C-E)**, consistent with the increased membrane trafficking shown in **Fig. 5A**. For WT, 24-hour treatment with E-4031 reduced currents (**Fig 5D**), indicating that the drug does not improve WT trafficking and the washout of drug is incomplete.

**Figure 5:**
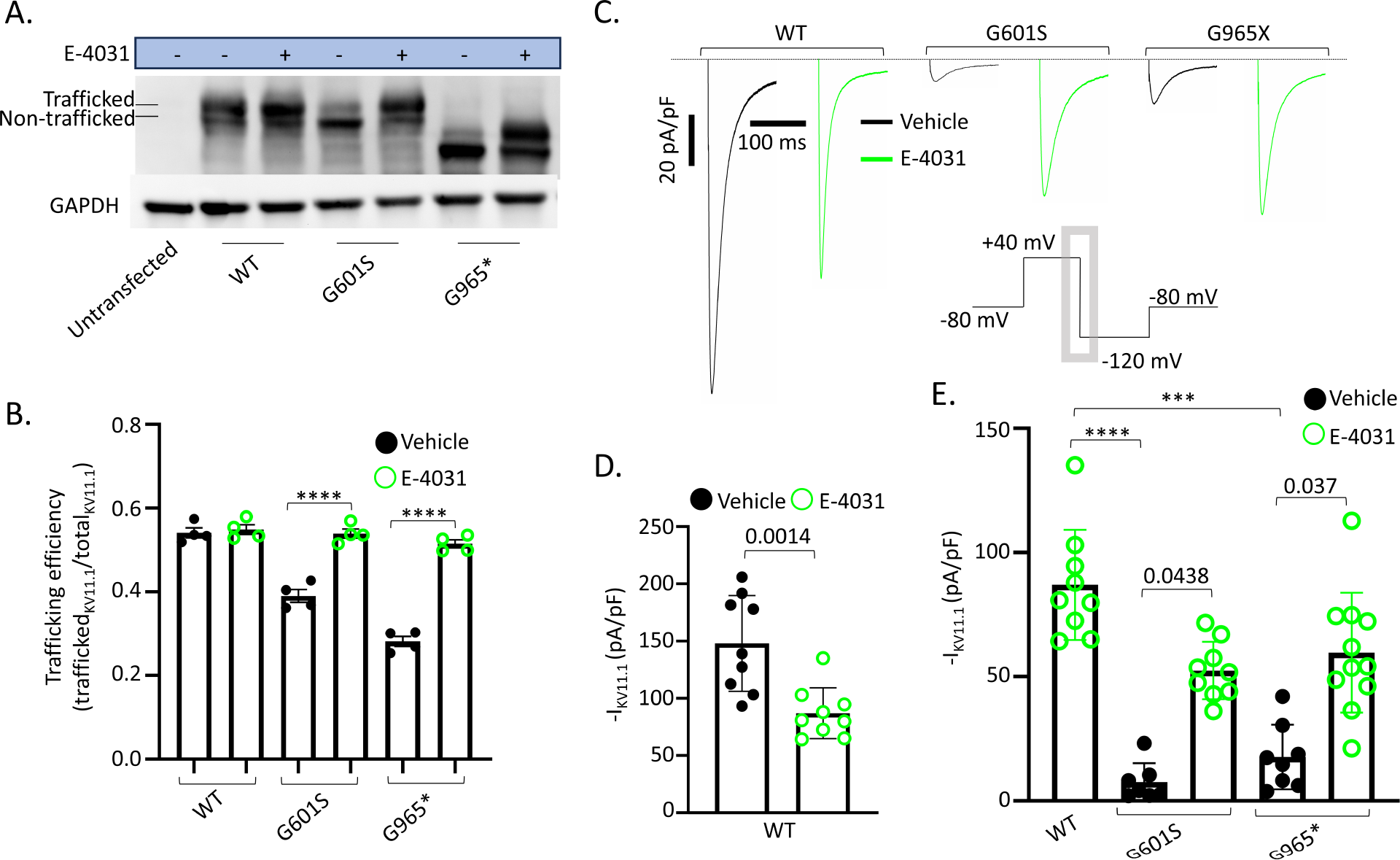
Impact of E-4031 on K_V_11.1 trafficking and function. **(A)** Western blot depicting K_V_11.1 and variant protein expression in HEK293 cells in the absence or presence of E-4031 for 24-hours. The truncated G965* has a lower molecular weight of ∼100 kD compared to 135 kD of full length Kv11.1. **(B)** Bar graph illustrating K_V_11.1 trafficking efficiency quantified by the ratio of trafficked 155 kD glycosylated protein to total K_V_11.1 protein, including untrafficked 135 kD K_V_11.1 protein. Statistical significance was calculated using one-way ANOVA with post-hoc Tukey’s multiple comparisons test and p values are depicted by **** < 0.0001. **(C)** Representative K_V_11.1 current traces at -120 mV step from voltage-clamp experiments after 24-hour treatment and washout. Inset: Voltage-clamp protocol. K_V_11.1 current was quantified as the peak inward current upon repolarization to -120 mV (gray box). **(D)** Bar graph comparing WT K_V_11.1 currents after 24-hour treatment and washout of vehicle or E-4031. Data analyzed with unpaired student’s t-test. **(E)** Bar graphs comparing WT K_V_11.1 current and two trafficking-deficient variants after 24-hour treatment with washout. All experiments follow a 24-hour treatment with vehicle (0.1% DMSO) or E-4031 (10 µM). Patch clamp experiments included a 30-minute washout to remove residual E-4031. Data are presented as mean ± SD. Statistical significance was calculated by Kruskal-Wallis with post-hoc Dunn’s multiple comparisons test and p values are depicted by **** < 0.0001, *** = 0.0001.

To date, little is known about the underlying mechanism of how E-4031 increases K_V_11.1 variant membrane trafficking and function. Therefore, we used our comparative interactome dataset for the two trafficking-deficient K_V_11.1 variants to evaluate which interactions were changed by E-4031 treatment. We found disparate responses for two variants. Globally, E-4031 altered protein interactions in the truncated variant G965* more than G601S (**Fig. 6A**). There was a 22% reduction in mean log_2_ fold change/WT in the G965* variant compared to only 7.6% reduction in the G601S variant (log_2_ fold change 1.69 to 1.56 G601S and 1.04 to 0.81 G965*). DAVID pathway analysis suggested that proteasomal proteins were significantly enriched in our dataset of K_V_11.1 interactors (**Fig 3B**). Thus, we scrutinized manually curated gene ontology terms related to proteasomal degradation from our data. Our data suggest that protein interactions with structural components of the proteasome (PSM) were significantly reduced in both variants treated with E-4031 (**Fig. 6B&C**). Manually annotated GO terms showed several individual proteins that had significant changes after E-4031 treatment in either variant (**Fig. S4**). Observing the top five protein interactions with the greatest change after E-4031 from 10 separate GO terms highlighted some key interactions in different pathways (**Fig. 6D**). Together these data suggest several interactions significantly altered after E-4031 treatment.

**Figure 6:**
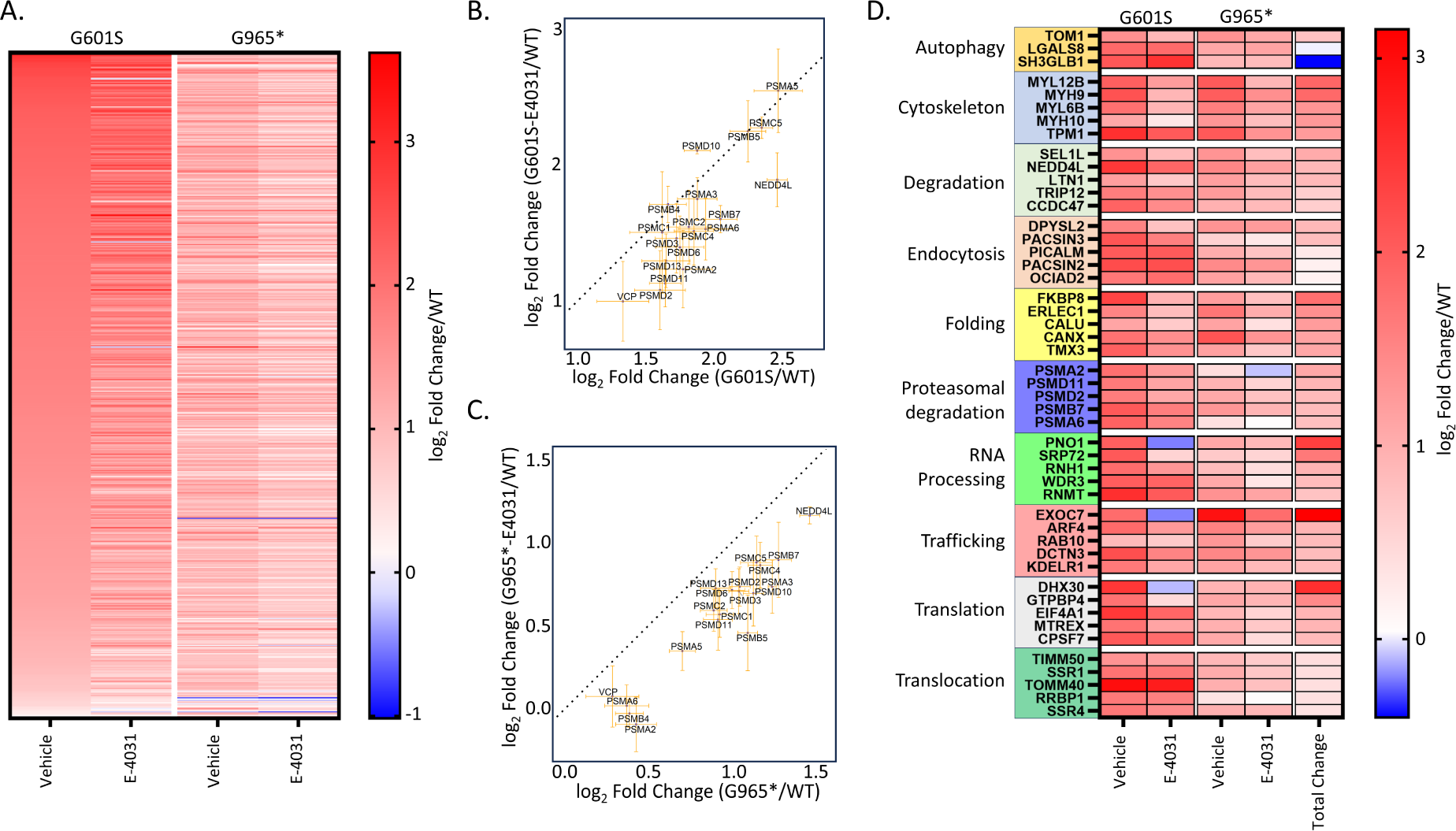
Impact of chaperone treatment on protein interactions in trafficking-deficient K_V_11.1 variants. **(A)** Heatmap illustrating 573 enriched protein interactions in trafficking deficient variants G601S and G965*, with or without 24-hour E-4031 treatment. **(B&C)** Correlation of proteasomal degradation protein interactions for **(B)** G601S and **(C)** G965* in absence (x-axis) or presence (y-axis) of E-4031 (all log_2_-fold changes normalized to WT with DMSO). The black dotted line indicates the normal line with a slope of 1 and intercept at the origin. The orange error bars from each individual protein represent the SEM for vehicle (0.1% DMSO, horizontal) and E-4031 (10 µM, vertical). **(D)** Heatmap showing the top five interactions from 10 manually annotated GO terms sorted by largest difference in log_2_ fold change/WT in absence or presence of E-4031 for 24 hours. Only three proteins responsible for autophagy were enriched for K_V_11.1 interaction in our dataset. Cells were treated with either vehicle (0.1% DMSO) or E-4031 (10 µ M) for 24-hours prior to experiments.

The following individual interactors were significantly reduced in both variants after E-4031 treatment: part of the exocyst complex responsible for docking exocytic vesicles with fusion sites in the plasma membrane (*EXOC7),* the ER-associated degradation protein sel-1 homolog 1 (*SEL1L*), the E3 ubiquitin ligase listerin (*LTN1*) involved in ribosomal quality control, and lectin based folding chaperones (*CANX* and *ERLEC)*. Interactions reduced after E-4031 treatment specific to the G601S variant included the folding chaperone peptidyl-prolyl cis-trans isomerase (*FKBP8*), the BCL2-associated athanogene 6 (*BAG6*), and a subunit of the multipass translocon (*CCDC47*). Interactions reduced by E-4031 treatment specific to the G965* variant included small GTPases involved in membrane trafficking and vesicle transport (*RAB10* and *RAB9A*) and a molecular chaperone in the BAG family (*BAG3*). Misfolded K_V_11.1 protein, unlike misfolded CFTR, has not been identified to aggregate *in vitro*. However, the interaction with *BAG6,* mainly in the full-length G601S variant, and formation of aggregates observed in purified K_V_11.1 protein indicate the potential aggregation of misfolded K_V_11.1.^6,37,38^ We observed many proteins responsible for clathrin-dependent or clathrin-independent endocytosis that remained unchanged after E-4031 treatment (**Fig. S5**). These experiments highlighted the proteostasis network associated with K_V_11.1 and indicate that specific variants may exhibit differential responses to pharmacological treatment. Here, we have shown the remodeling of the proteostasis network is larger in the truncated variant (G965*) even with lower interactions before treatment. This understanding is crucial for guiding future research and developing therapeutic approaches.

## Discussion

This study is the first to investigate the interactomes of trafficking-deficient K_V_11.1 variants and compare them to WT K_V_11.1 using AP-MS coupled with TMT labeling. Through this approach, we identified several proteins already known to interact with K_V_11.1, such as the E3 ubiquitin ligase (*NEDD4L)*, the four-and-a-half LIM domains protein 2 (*FHL2)*, the lectin-based folding chaperone calnexin (*CANX)*, Caveolin-1 (*CAV1*), and the 14-3-3ε protein (*YWHAE*).^32,33,39–42^ The truncated variant interaction with 14-3-3ε was reduced compared to either WT or the full-length K_V_11.1-G601S variant, consistent with previous work that suggests 14-3-3ε interacts with the C-terminal domain.^40^ K_V_11.1, a glyocoprotein, undergoes N-linked glycosylation at the N598 residue, and mutations at this site were shown to influence channel stability.^43^ We identified several proteins involved in glycosylation or glycoprotein folding: protein glucosyltransferases (*DDOST* and *RPN2*), lectin based chaperones (*ERLEC* and *CANX*), and glucosidases (*MOGS* and *GANAB*). We also identified protein interactions that have not been previously reported such as: the estradiol 17-beta hydoxysteroid dehydrogenase (*HSD17B12*), the BCL2-associated athanogene (*BAG6*), and mitochondrial membrane-associated or solute carrier proteins (TOMM40, TOMM70, TOMM22, TIMM50, TRAP1, SLC25A10, SLC25A13, SLC25A24, SLC25A3). Several mitochondrial proteins were found to interact with gamma-aminobutyric acid type A (GABA_A_) channels as well and warrant further exploration into possible functional roles on K_V_11.1 biogenesis.^31^ The interaction with *HSD17B12,* which converts estrone to estradiol could be particularly significant given that female sex hormones increase the risk of LQTS related arrhythmias.^44,45^ Several other 17-beta hydroxysteroid dehydrogenases localize in the ER membrane, and the catalytic site of *HSD17B12* was found facing the ER lumen.^46^ Thus this interaction could have early effects on K_V_11.1 folding within the ER.

E-4031 improves trafficking efficiency of K_V_11.1 variants and our data show an associated decrease in protein interactions after treatment in K_V_11.1 variants. While the direct mechanism of E-4031 remains unidentified, we discovered numerous protein interactions that are altered with E-4031 treatment and could be therapeutically targeted. Several studies have demonstrated that knockdown via siRNA or overexpression of folding proteins can improve the trafficking and function of K_V_11.1 variants.^47–51^ Experiments of this nature hold significance not only for K_V_11.1 but also in the context of cystic fibrosis, where research has revealed several protein interactions that could be modulated to enhance CFTR trafficking.^14,15^ Similarly, other ion channel trafficking disorders present promising protein interaction targets.^52–54^ Continued research in these domains may not only yield novel treatments for K_V_11.1-associated conditions but also benefit a broader spectrum of ion channel trafficking disorders.

An important limitation of this study was the use of HEK293 cells, which differ considerably from native cardiomyocytes and do not express all proteins and channels present in a cardiomyocyte. Nonetheless, HEK293 cells are a commonly-employed model cell line to study the secretory pathway, and they have been able to recapitulate many of the native proteostasis interactions of other membrane proteins.^14,31,55^ To overcome this limitation, future investigations could use human induced pluripotent stem cell cardiomyocytes to identify protein interactions in a more native-like cell type. Another limitation of our study is that known interactors did not show significant enrichment in K_V_11.1 pulldown samples, such as the heat-shock protein (Hsp70) family. We identified several Hsp70 family member proteins (HSPA-1A/4/4L/5/6/8) but these were not enriched compared to untransfected cells.^34,45^ This suggests that our mass spectrometry samples may contain false negatives due to substantial protein abundance in the untransfected mock IP samples, as has been seen in the Contaminant Repository for Affinity Purification (CRAPome).^56^

Our data suggest that E-4031 does not improve the trafficking efficiency of WT, implying a potential saturation in the channel’s trafficking rate. Furthermore, E-4031 does not appear to reduce degradation of the WT channel, nor modify specific protein interactions responsible for internalization of K_V_11.1 variants. This observation is supported by the lack of change in protein interactions associated with clathrin mediated internalization (*SPIRE1* and *PICALM)* and non-clathrin mediated endocytosis (*CAV1, CAVIN1,* and *SPTBN2*) **(Fig S5)**. As such, E-4031 may predominantly promote the forward trafficking of K_V_11.1 variants, exerting minimal influence on their endocytosis. The plasma surface half-life of K_V_11.1 variants has been shown to be shorter than WT, suggesting an alternative therapeutic strategy targeting plasma membrane stabilization could be explored.^57,58^ Clathrin-mediated endocytosis has not been found as a major mechanism of K_V_11.1 internalization, but caveolin and NEDD4L are thought to mediate internalization during extracellular potassium depletion.^39^ We identified several E3 ubiquitin ligases that were not previously known to interact with K_V_11.1, including *UBE2C*, *LTN1*, *ITCH*, *TRIP12*, *UBR3*, *MIB1*, and *CUL3*. This evidence suggests that E3 ubiquitin ligases could play a large role in channel turnover, particularly at the plasma membrane.

Our findings indicate that removal of the ER retention motif and C-terminal domain could alter protein interactions responsible for trafficking of K_V_11.1 channels. It is plausible that the ER retention motif, RGR (residues 1005-1007) could be therapeutically targeted to increase trafficking. Others reported that truncated K_V_11.1 variants with the ER retention domain intact have no functional current, which was correctable upon deletion or treatment with a peptide spanning the domain.^59^ In our study, we found that the truncated variant displayed smaller interactome differences compared to WT protein than the full-length variant. Our dataset highlighted the proteasomal degradation pathway, suggesting that K_V_11.1 protein and variants are primarily degraded by the proteasome as opposed to lysosomal degradation. Many of the protein interactions linked to proteasomal degradation were decreased after E-4031 treatment, further supporting this pathway could be a target for intervention.

In conclusion, we have for the first time defined the protein interactomes of K_V_11.1, two of its trafficking-deficient variants, and the effect of a pharmacological chaperone (E-4031) on the protein interaction network. Future research will have to elucidate functional implications of specific protein-protein interactions identified in this study and clarify which interactions could become relevant targets for therapeutic strategies in LQTS.

## Acknowledgments

Special thanks to the laboratory of Dan Roden for supply of HEK293 cells. Special thanks to Hayes McDonald for guidance on coimmunoprecipitation procedures.

## Funding info

This work was supported in part by the National Institutes of Health (NIH) grants [NHLBI R35 HL144980] (B.C.K., T.D.), [NIGMS R35 GM133552] (L.P., L.B.), [NHLBI R01 HL167046] (L.P.), [T32 – 5T32GM007569-44] (B.C.K., C.L.E.), [NHLBI F31 HL162483] (E.F.M); and the Leducq Foundation grant [18CVD05] (B.C.K.).

## Author contributions

Research design – Egly, Plate, Knollmann

Experimental work: Egly, Do, Barny

Writing – original draft preparation: Egly, Plate, Knollmann

Writing – review and editing: Egly, Plate, Knollmann, McDonald, Do, Barny

Contributed new reagents of analytic tools: Plate, Barny, McDonald

Data Analysis: Egly, Barny

## References

1. Schwartz PJ, Stramba-Badiale M, Crotti L, Pedrazzini M, Besana A, Bosi G, Gabbarini F, Goulene K, Insolia R, Mannarino S, et al. Prevalence of the congenital long-QT syndrome. Circulation. 2009;120:1761–1767. doi: 10.1161/CIRCULATIONAHA.109.863209

2. Wilde AAM, Amin AS, Postema PG. Diagnosis, management and therapeutic strategies for congenital long QT syndrome. Heart. 2022;108:332–338. doi: 10.1136/heartjnl-2020-318259

3. Bos JM, Crotti L, Rohatgi RK, Castelletti S, Dagradi F, Schwartz PJ, Ackerman MJ. Mexiletine Shortens the QT Interval in Patients With Potassium Channel-Mediated Type 2 Long QT Syndrome. Circ Arrhythm Electrophysiol. 2019;12:e007280. doi: 10.1161/CIRCEP.118.007280

4. Anderson CL, Kuzmicki CE, Childs RR, Hintz CJ, Delisle BP, January CT. Large-scale mutational analysis of Kv11.1 reveals molecular insights into type 2 long QT syndrome. Nat Commun. 2014;5:5535. doi: 10.1038/ncomms6535

5. Sanguinetti MC, Tristani-Firouzi M. hERG potassium channels and cardiac arrhythmia. Nature. 2006;440:463–469. doi: 10.1038/nature04710

6. Wang W, MacKinnon R. Cryo-EM Structure of the Open Human Ether-à-go-go-Related K. Cell. 2017;169:422–430.e410. doi: 10.1016/j.cell.2017.03.048

7. Smith JL, Reloj AR, Nataraj PS, Bartos DC, Schroder EA, Moss AJ, Ohno S, Horie M, Anderson CL, January CT, et al. Pharmacological correction of long QT-linked mutations in KCNH2 (hERG) increases the trafficking of Kv11.1 channels stored in the transitional endoplasmic reticulum. Am J Physiol Cell Physiol. 2013;305:C919–930. doi: 10.1152/ajpcell.00406.2012

8. Zhou Z, Gong Q, January CT. Correction of defective protein trafficking of a mutant HERG potassium channel in human long QT syndrome. Pharmacological and temperature effects. J Biol Chem. 1999;274:31123–31126. doi: 10.1074/jbc.274.44.31123

9. Gong Q, Jones MA, Zhou Z. Mechanisms of pharmacological rescue of trafficking-defective hERG mutant channels in human long QT syndrome. J Biol Chem. 2006;281:4069–4074. doi: 10.1074/jbc.M511765200

10. Ficker E, Obejero-Paz CA, Zhao S, Brown AM. The binding site for channel blockers that rescue misprocessed human long QT syndrome type 2 ether-a-gogo-related gene (HERG) mutations. J Biol Chem. 2002;277:4989–4998. doi: 10.1074/jbc.M107345200

11. Hartl FU, Bracher A, Hayer-Hartl M. Molecular chaperones in protein folding and proteostasis. Nature. 2011;475:324–332. doi: 10.1038/nature10317

12. Scheufler C, Brinker A, Bourenkov G, Pegoraro S, Moroder L, Bartunik H, Hartl FU, Moarefi I. Structure of TPR domain-peptide complexes: critical elements in the assembly of the Hsp70-Hsp90 multichaperone machine. Cell. 2000;101:199–210. doi: 10.1016/s0092-8674(00)80830-2

13. Betegon M, Brodsky JL. Unlocking the door for ERAD. Nat Cell Biol. 2020;22:263–265. doi: 10.1038/s41556-020-0476-1

14. McDonald EF, Sabusap CMP, Kim M, Plate L. Distinct proteostasis states drive pharmacologic chaperone susceptibility for cystic fibrosis transmembrane conductance regulator misfolding mutants. Mol Biol Cell. 2022;33:ar62. doi: 10.1091/mbc.E21-11-0578

15. Pankow S, Bamberger C, Calzolari D, Martínez-Bartolomé S, Lavallée-Adam M, Balch WE, Yates JR. ΔF508 CFTR interactome remodelling promotes rescue of cystic fibrosis. Nature. 2015;528:510–516. doi: 10.1038/nature15729

16. Jones DK, Johnson AC, Roti Roti EC, Liu F, Uelmen R, Ayers RA, Baczko I, Tester DJ, Ackerman MJ, Trudeau MC, et al. Localization and functional consequences of a direct interaction between TRIOBP-1 and hERG proteins in the heart. J Cell Sci. 2018;131. doi: 10.1242/jcs.206730

17. Ledford HA, Ren L, Thai PN, Park S, Timofeyev V, Sirish P, Xu W, Emigh AM, Priest JR, Perez MV, et al. Disruption of protein quality control of the human ether-à-go-go related gene K(+) channel results in profound long QT syndrome. Heart Rhythm. 2022;19:281–292. doi: 10.1016/j.hrthm.2021.10.005

18. Roder K, Kabakov A, Moshal KS, Murphy KR, Xie A, Dudley S, Turan NN, Lu Y, MacRae CA, Koren G. Trafficking of the human ether-a-go-go-related gene (hERG) potassium channel is regulated by the ubiquitin ligase rififylin (RFFL). J Biol Chem. 2019;294:351–360. doi: 10.1074/jbc.RA118.003852

19. Iwai C, Li P, Kurata Y, Hoshikawa Y, Morikawa K, Maharani N, Higaki K, Sasano T, Notsu T, Ishido Y, et al. Hsp90 prevents interaction between CHIP and HERG proteins to facilitate maturation of wild-type and mutant HERG proteins. Cardiovasc Res. 2013;100:520–528. doi: 10.1093/cvr/cvt200

20. Kang Y, Guo J, Yang T, Li W, Zhang S. Regulation of the human ether-a-go-go-related gene (hERG) potassium channel by Nedd4 family interacting proteins (Ndfips). Biochem J. 2015;472:71–82. doi: 10.1042/BJ20141282

21. Marinko JT, Wright MT, Schlebach JP, Clowes KR, Heintzman DR, Plate L, Sanders CR. Glycosylation limits forward trafficking of the tetraspan membrane protein PMP22. J Biol Chem. 2021;296:100719. doi: 10.1016/j.jbc.2021.100719

22. Wright MT, Kouba L, Plate L. Thyroglobulin Interactome Profiling Defines Altered Proteostasis Topology Associated With Thyroid Dyshormonogenesis. Mol Cell Proteomics. 2021;20:100008. doi: 10.1074/mcp.RA120.002168

23. Egly CL, Blackwell DJ, Schmeckpeper J, Delisle BP, Weaver CD, Knollmann BC. A High-Throughput Screening Assay to Identify Drugs that Can Treat Long QT Syndrome Caused by Trafficking-Deficient KV11.1 (hERG) Variants. Mol Pharmacol. 2022;101:236–245. doi: 10.1124/molpharm.121.000421

24. Pankow S, Bamberger C, Calzolari D, Bamberger A, Yates JR, 3rd. Deep interactome profiling of membrane proteins by co-interacting protein identification technology. Nat Protoc. 2016;11:2515–2528. doi: 10.1038/nprot.2016.140

25. DeCaprio J, Kohl TO. Cross-Linking Antibodies to Beads Using Dimethyl Pimelimidate (DMP). Cold Spring Harb Protoc. 2019;2019. doi: 10.1101/pdb.prot098624

26. Huang da W, Sherman BT, Lempicki RA. Systematic and integrative analysis of large gene lists using DAVID bioinformatics resources. Nat Protoc. 2009;4:44–57. doi: 10.1038/nprot.2008.211

27. Huang da W, Sherman BT, Lempicki RA. Bioinformatics enrichment tools: paths toward the comprehensive functional analysis of large gene lists. Nucleic Acids Res. 2009;37:1–13. doi: 10.1093/nar/gkn923

28. Benjamini Y, Hochberg Y. On the Adaptive Control of the False Discovery Rate in Multiple Testing with Independent Statistics. Journal of Educational and Behavioral Studies. 2000;25:60–83.

29. Furutani M, Trudeau MC, Hagiwara N, Seki A, Gong Q, Zhou Z, Imamura S, Nagashima H, Kasanuki H, Takao A, et al. Novel mechanism associated with an inherited cardiac arrhythmia: defective protein trafficking by the mutant HERG (G601S) potassium channel. Circulation. 1999;99:2290–2294. doi: 10.1161/01.cir.99.17.2290

30. Kim M, McDonald EF, Sabusap CMP, Timalsina B, Joshi D, Hong JS, Rab A, Sorscher EJ, Plate L. Elexacaftor/VX-445-mediated CFTR interactome remodeling reveals differential correction driven by mutation-specific translational dynamics. J Biol Chem. 2023;299:105242. doi: 10.1016/j.jbc.2023.105242

31. Wang YJ, Di XJ, Mu TW. Quantitative interactome proteomics identifies a proteostasis network for GABA(A) receptors. J Biol Chem. 2022;298:102423. doi: 10.1016/j.jbc.2022.102423

32. Wu Y, Huang X, Zheng Z, Yang X, Ba Y, Lian J. Role and mechanism of chaperones calreticulin and ERP57 in restoring trafficking to mutant HERG-A561V protein. Int J Mol Med. 2021;48. doi: 10.3892/ijmm.2021.4992

33. Kagan A, Melman YF, Krumerman A, McDonald TV. 14-3-3 amplifies and prolongs adrenergic stimulation of HERG K+ channel activity. Embo j. 2002;21:1889–1898. doi: 10.1093/emboj/21.8.1889

34. Kagan A, McDonald TV. Dynamic control of hERG/I(Kr) by PKA-mediated interactions with 14-3-3. Novartis Found Symp. 2005;266:75–89; discussion 89-99.

35. Lu J, Robinson JM, Edwards D, Deutsch C. T1-T1 interactions occur in ER membranes while nascent Kv peptides are still attached to ribosomes. Biochemistry. 2001;40:10934–10946. doi: 10.1021/bi010763e

36. Li K, Jiang Q, Bai X, Yang YF, Ruan MY, Cai SQ. Tetrameric Assembly of K(+) Channels Requires ER-Located Chaperone Proteins. Mol Cell. 2017;65:52–65. doi: 10.1016/j.molcel.2016.10.027

37. Kleizen B, de Mattos E, Papaioannou O, Monti M, Tartaglia GG, van der Sluijs P, Braakman I. Transmembrane Helices 7 and 8 Confer Aggregation Sensitivity to the Cystic Fibrosis Transmembrane Conductance Regulator. Int J Mol Sci. 2023;24. doi: 10.3390/ijms242115741

38. Kawahara H, Minami R, Yokota N. BAG6/BAT3: emerging roles in quality control for nascent polypeptides. J Biochem. 2013;153:147–160. doi: 10.1093/jb/mvs149

39. Guo J, Wang T, Li X, Shallow H, Yang T, Li W, Xu J, Fridman MD, Yang X, Zhang S. Cell surface expression of human ether-a-go-go-related gene (hERG) channels is regulated by caveolin-3 protein via the ubiquitin ligase Nedd4-2. J Biol Chem. 2012;287:33132–33141. doi: 10.1074/jbc.M112.389643

40. Choe CU, Schulze-Bahr E, Neu A, Xu J, Zhu ZI, Sauter K, Bähring R, Priori S, Guicheney P, Mönnig G, et al. C-terminal HERG (LQT2) mutations disrupt IKr channel regulation through 14-3-3epsilon. Hum Mol Genet. 2006;15:2888–2902. doi: 10.1093/hmg/ddl230

41. Ma Q, Yu H, Lin J, Sun Y, Shen X, Ren L. Screening for cardiac HERG potassium channel interacting proteins using the yeast two-hybrid technique. Cell Biol Int. 2014;38:239–245. doi: 10.1002/cbin.10196

42. Maurya S, Mills RW, Kahnert K, Chiang DY, Bertoli G, Lundegaard PR, Duran MP-H, Zhang M, Rothenberg E, George AL, et al. Outlining cardiac ion channel protein interactors and their signature in the human electrocardiogram.

43. Gong Q, Anderson CL, January CT, Zhou Z. Role of glycosylation in cell surface expression and stability of HERG potassium channels. Am J Physiol Heart Circ Physiol. 2002;283:H77–84. doi: 10.1152/ajpheart.00008.2002

44. Yang PC, Perissinotti LL, López-Redondo F, Wang Y, DeMarco KR, Jeng MT, Vorobyov I, Harvey RD, Kurokawa J, Noskov SY, et al. A multiscale computational modelling approach predicts mechanisms of female sex risk in the setting of arousal-induced arrhythmias. J Physiol. 2017;595:4695–4723. doi: 10.1113/jp273142

45. Bjelic M, Zareba W, Peterson DR, Younis A, Aktas MK, Huang DT, Rosero S, Cutter K, McNitt S, Xia X, et al. Sex hormones and repolarization dynamics during the menstrual cycle in women with congenital long QT syndrome. Heart Rhythm. 2022;19:1532–1540. doi: 10.1016/j.hrthm.2022.04.029

46. Tsachaki M, Odermatt A. Subcellular localization and membrane topology of 17β-hydroxysteroid dehydrogenases. Mol Cell Endocrinol. 2019;489:98–106. doi: 10.1016/j.mce.2018.07.003

47. Apaja PM, Foo B, Okiyoneda T, Valinsky WC, Barriere H, Atanasiu R, Ficker E, Lukacs GL, Shrier A. Ubiquitination-dependent quality control of hERG K+ channel with acquired and inherited conformational defect at the plasma membrane. Mol Biol Cell. 2013;24:3787–3804. doi: 10.1091/mbc.E13-07-0417

48. Ficker E, Dennis AT, Wang L, Brown AM. Role of the cytosolic chaperones Hsp70 and Hsp90 in maturation of the cardiac potassium channel HERG. Circ Res. 2003;92:e87–100. doi: 10.1161/01.RES.0000079028.31393.15

49. Walker VE, Atanasiu R, Lam H, Shrier A. Co-chaperone FKBP38 promotes HERG trafficking. J Biol Chem. 2007;282:23509–23516. doi: 10.1074/jbc.M701006200

50. Walker VE, Wong MJ, Atanasiu R, Hantouche C, Young JC, Shrier A. Hsp40 chaperones promote degradation of the HERG potassium channel. J Biol Chem. 2010;285:3319–3329. doi: 10.1074/jbc.M109.024000

51. Hantouche C, Williamson B, Valinsky WC, Solomon J, Shrier A, Young JC. Bag1 Co-chaperone Promotes TRC8 E3 Ligase-dependent Degradation of Misfolded Human Ether a Go-Go-related Gene (hERG) Potassium Channels. J Biol Chem. 2017;292:2287–2300. doi: 10.1074/jbc.M116.752618

52. Di XJ, Han DY, Wang YJ, Chance MR, Mu TW. SAHA enhances Proteostasis of epilepsy-associated α1(A322D)β2γ2 GABA(A) receptors. Chem Biol. 2013;20:1456–1468. doi: 10.1016/j.chembiol.2013.09.020

53. Fu YL, Han DY, Wang YJ, Di XJ, Yu HB, Mu TW. Remodeling the endoplasmic reticulum proteostasis network restores proteostasis of pathogenic GABAA receptors. PLoS One. 2018;13:e0207948. doi: 10.1371/journal.pone.0207948

54. Mu TW, Ong DS, Wang YJ, Balch WE, Yates JR, 3rd, Segatori L, Kelly JW. Chemical and biological approaches synergize to ameliorate protein-folding diseases. Cell. 2008;134:769–781. doi: 10.1016/j.cell.2008.06.037

55. Geiger T, Wehner A, Schaab C, Cox J, Mann M. Comparative proteomic analysis of eleven common cell lines reveals ubiquitous but varying expression of most proteins. Mol Cell Proteomics. 2012;11:M111.014050. doi: 10.1074/mcp.M111.014050

56. Mellacheruvu D, Wright Z, Couzens AL, Lambert JP, St-Denis NA, Li T, Miteva YV, Hauri S, Sardiu ME, Low TY, et al. The CRAPome: a contaminant repository for affinity purification-mass spectrometry data. Nat Methods. 2013;10:730–736. doi: 10.1038/nmeth.2557

57. Kanner SA, Jain A, Colecraft HM. Development of a High-Throughput Flow Cytometry Assay to Monitor Defective Trafficking and Rescue of Long QT2 Mutant hERG Channels. Front Physiol. 2018;9:397. doi: 10.3389/fphys.2018.00397

58. Foo B, Barbier C, Guo K, Vasantharuban J, Lukacs GL, Shrier A. Mutation-specific peripheral and ER quality control of hERG channel cell-surface expression. Sci Rep. 2019;9:6066. doi: 10.1038/s41598-019-42331-6

59. Kupershmidt S, Yang T, Chanthaphaychith S, Wang Z, Towbin JA, Roden DM. Defective human Ether-à-go-go-related gene trafficking linked to an endoplasmic reticulum retention signal in the C terminus. J Biol Chem. 2002;277:27442–27448. doi: 10.1074/jbc.M112375200

