## Supplemental figures for "The proteostasis interactomes of trafficking-deficient K_V_11.1 variants associated with Long QT Syndrome and pharmacological chaperone rescue"

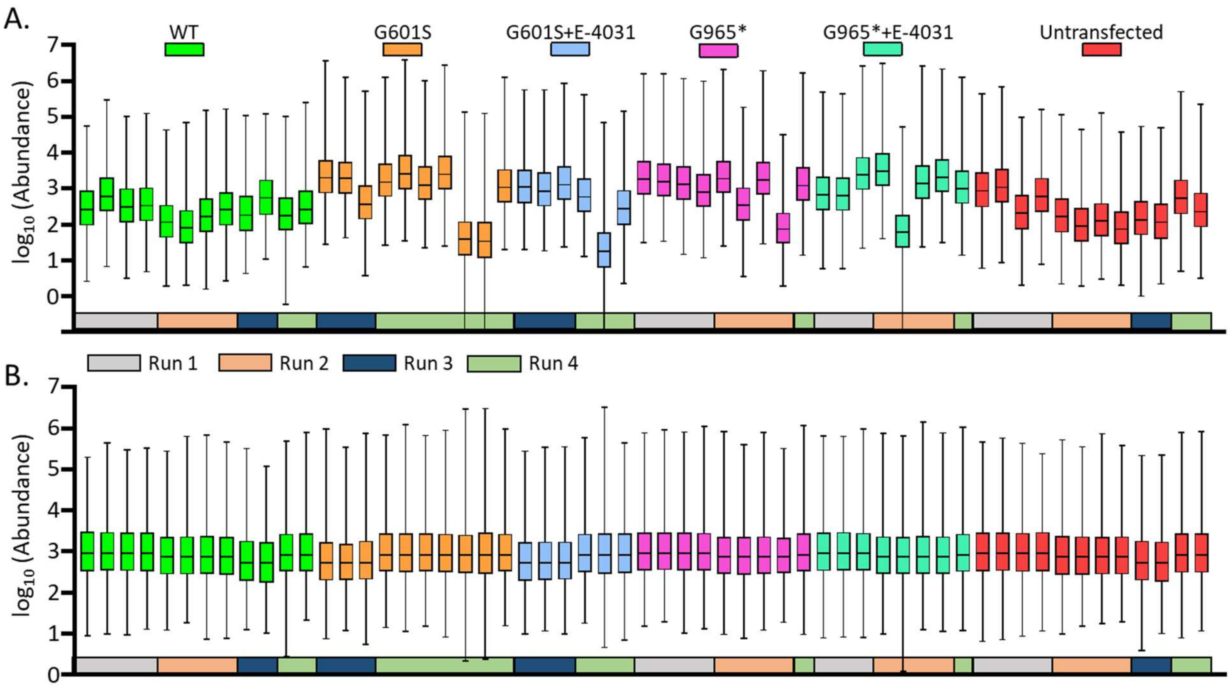


**Figure S1: Kv11.1 TMT intensity distribution and normalization. (A)** Box-and-whisker plot of $log_{10}$ TMT intensity abundance for all 4 mass spectrometry runs used in this study. Each cell line and treatment condition are grouped by the same color boxes with identity of the corresponding mass spectrometry run featured below in different colored boxes. **(B)** Box-and-whisker plot of $log_{10}$ TMT intensity abundance from A, but TMT abundance was median normalized for each mass spectrometry run.

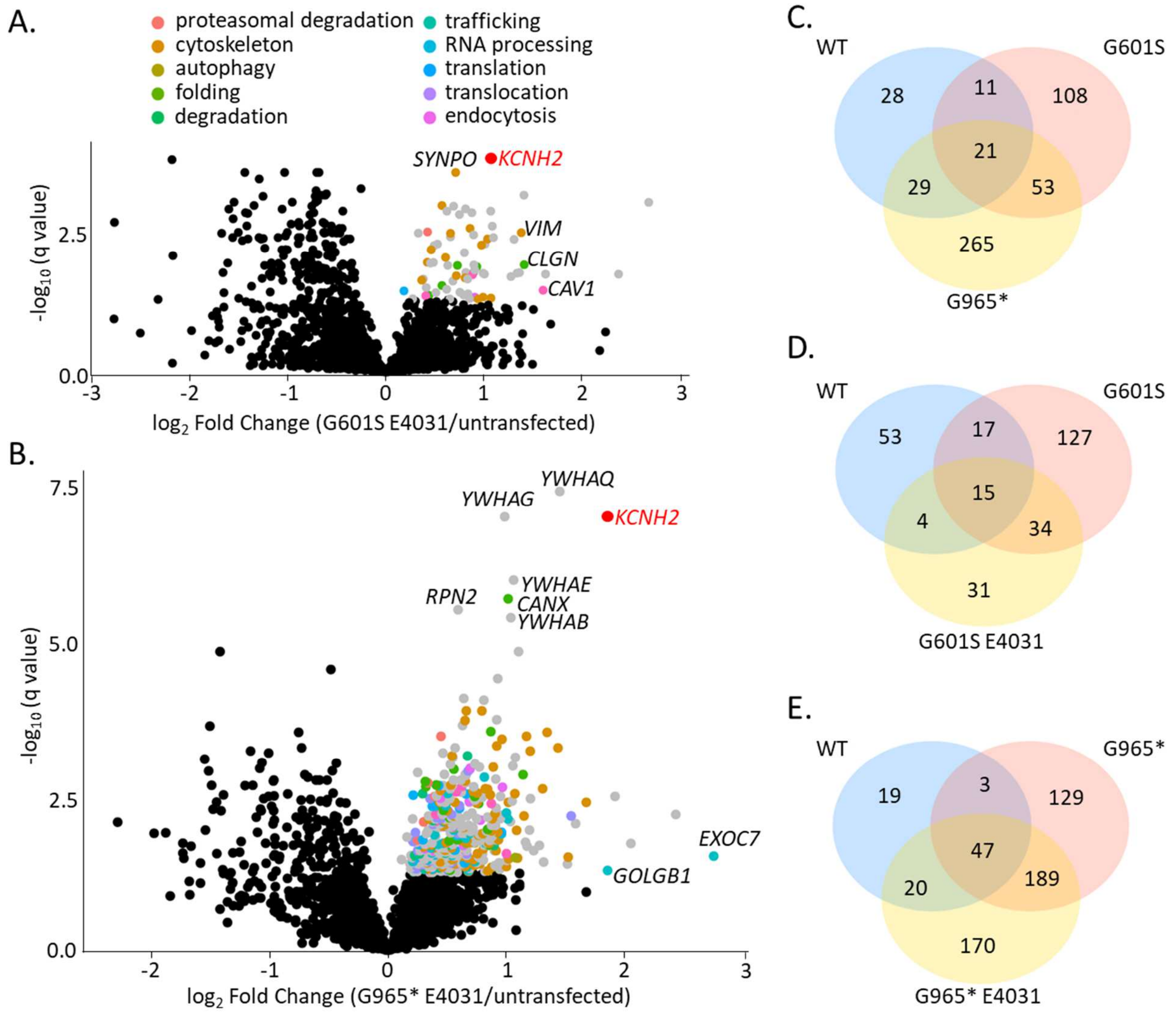


**Figure S2: Analysis of protein interactions in E-4031 treated Kv11.1 variants. (A&B)** Volcano plots showing identification of key interactors using multiplexed AP-MS proteomics for two variants **(A)** G601S and **(B)** G965* treated with E-4031 for 24-hours. These plots highlight significantly upregulated proteins in gray ($q < 0.05$ by multiple t-tests with false discovery rate correction using two-stage linear step-up of Benjamini, Krieger, and Yekutieli) compared to untransfected cell lines. Proteins are further categorized using manually curated gene ontology terms (N=10), color-coded for clarity. Key proteins such as Kv11.1 (*KCNH2,* red) and other genes are labeled. **(C)** Venn diagram showing overlap of Kv11.1 proteins significantly enriched in WT and variants. **(D-E)** Venn diagram showing overlap of protein interactions between WT and G601S **(D)** or G965* **(E)** and treated with vehicle (0.1% DMSO) or E-4031 (10 µM) for 24-hours.

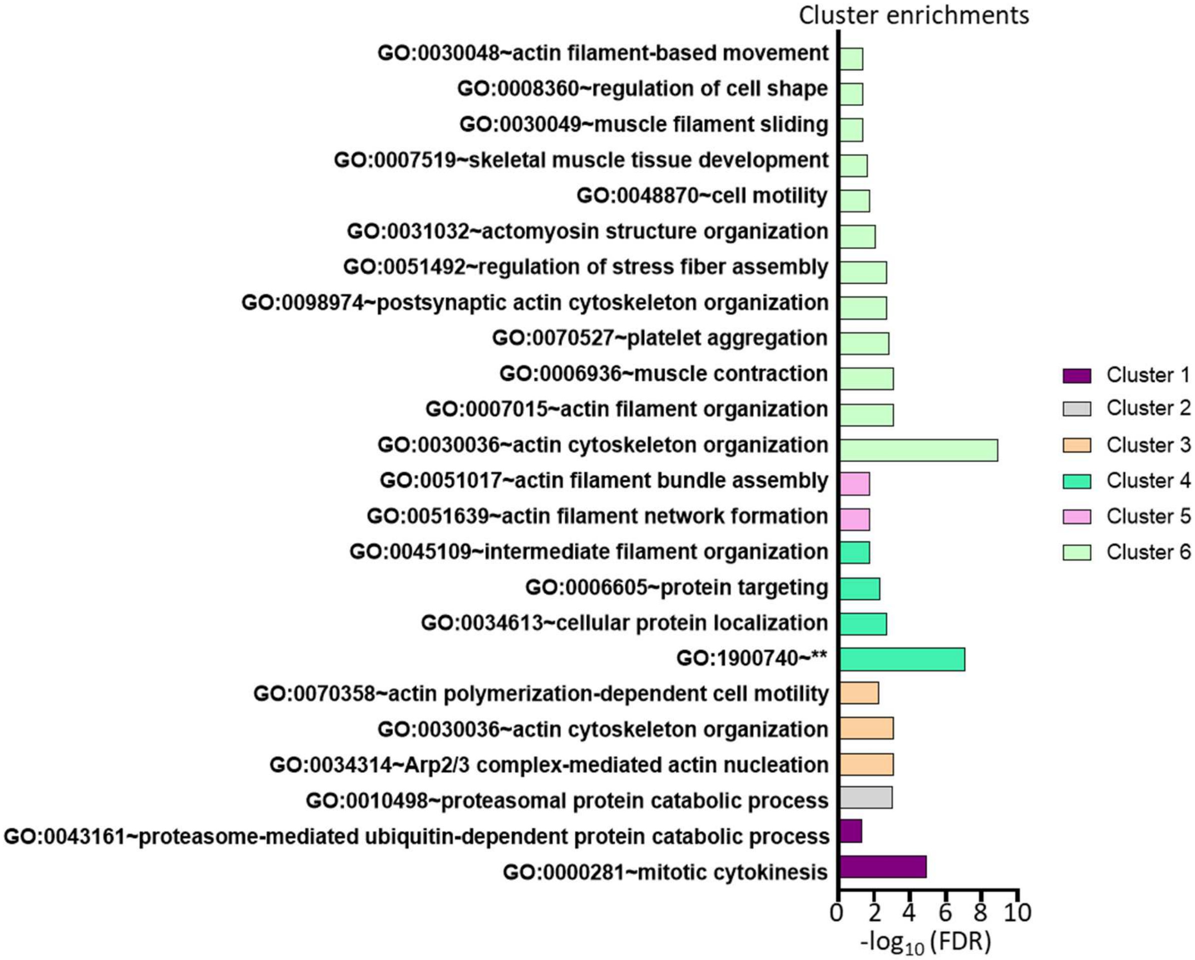


**Figure S3: K$_V$11.1 variant cluster enrichment with DAVID analysis.** Bar chart showing all significantly enriched GO terms based on hierarchical clustering of 573 protein interactors. Clusters are based on intensity similarity where wild-type K$_V$11.1 normalized abundances for the G601S and G965* variants were converted to a Euclidian distance matrix and clustered using Ward's minimum variance method. Statistical significance was determined using DAVID analysis by linear step up procedure using Benjamini Hochberg to control for false discovery rate ($q < 0.05$). ** = positive regulation of protein insertion into mitochondrial membrane involved in apoptotic signaling pathway.

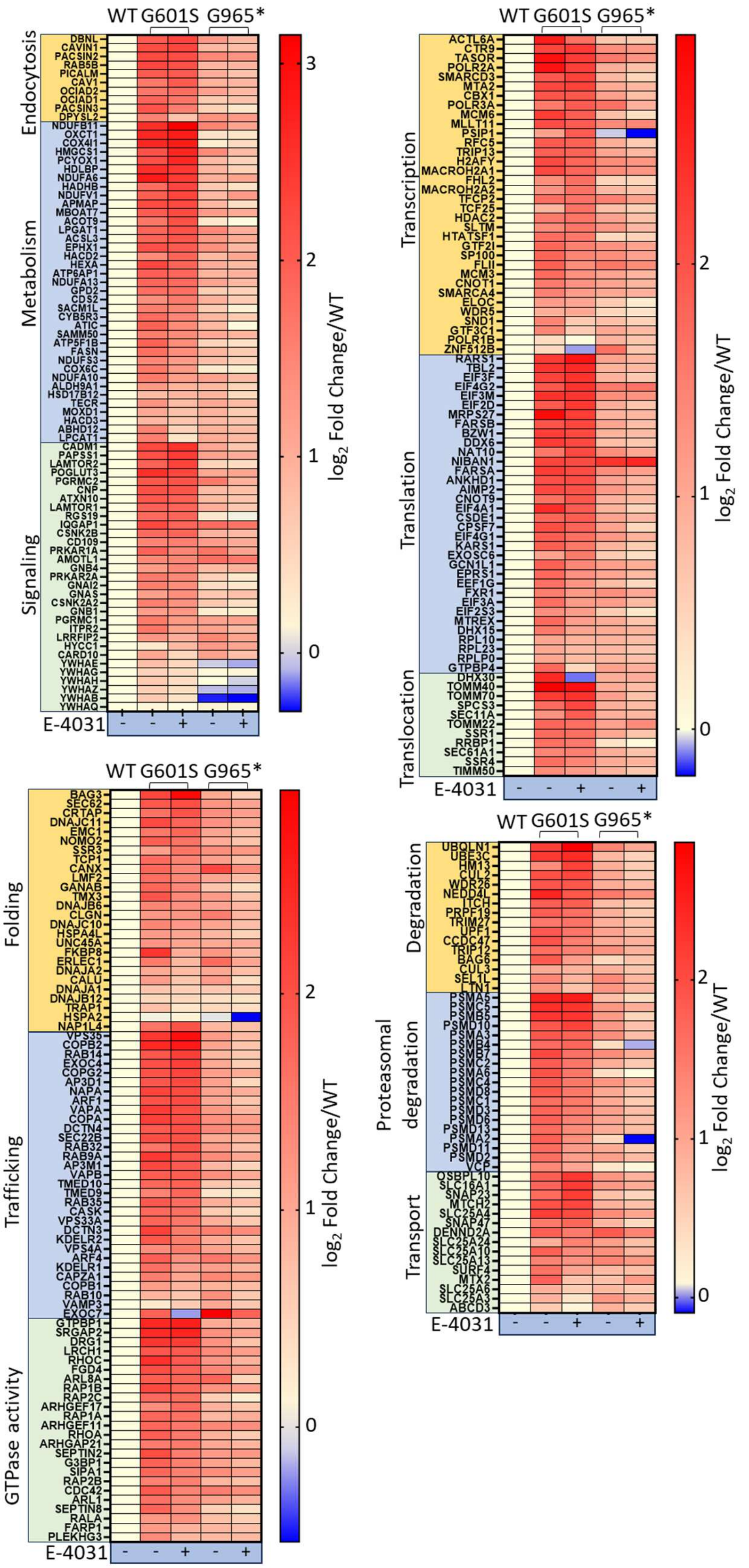


**Fig S4: Heatmaps of $K_V$11.1 interactors with annotated GO terms.** Heatmaps illustrating the protein interactions changes in $K_V$11.1 variants after 24-hour treatment with vehicle (0.1% DMSO) or E-4031 (10 μM). $Log_2$ fold change of protein abundance is scaled to wild-type and grouped into 12 separate manually annotated GO terms.

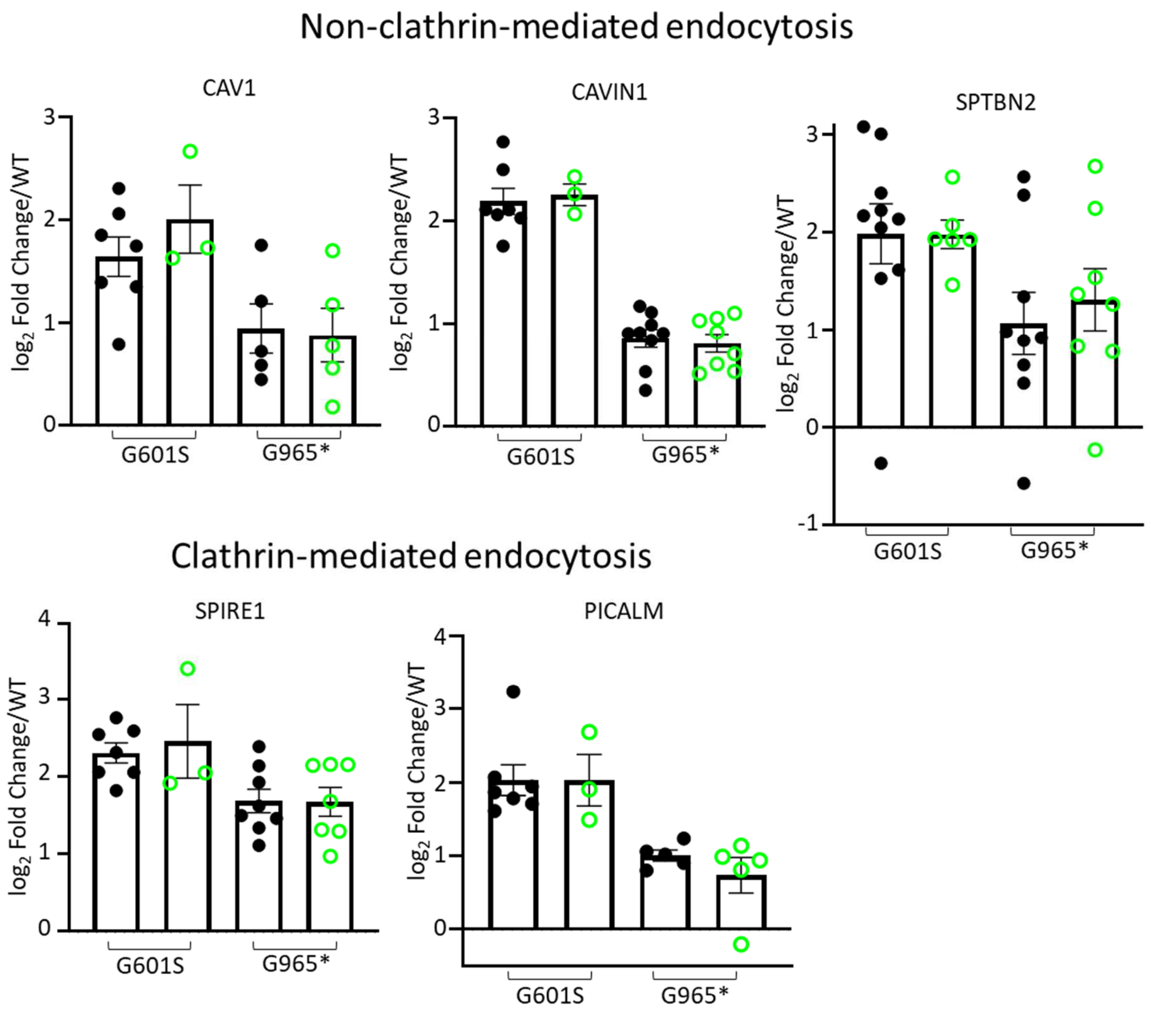


**Fig S5: No change in Kv11.1 endocytosis protein interactions with E-4031 treatment.** Bar charts showing $\log_2$ fold change for protein abundance normalized to WT for five key protein interactions responsible for endocytosis. Dots represent single replicates from mass spectrometry runs for each group. Either variant was treated with vehicle (DMSO 0.1%) or E-4031 (10 μM) for 24-hours prior to experiments.
